# Topoisomerase 1 inhibition therapy protects against SARS-CoV-2-induced inflammation and death in animal models

**DOI:** 10.1101/2020.12.01.404483

**Authors:** Jessica Sook Yuin Ho, Bobo Wing-Yee Mok, Laura Campisi, Tristan Jordan, Soner Yildiz, Sreeja Parameswaran, Joseph A Wayman, Natasha N Gaudreault, David A Meekins, Sabarish V. Indran, Igor Morozov, Jessie D Trujillo, Yesai S Fstkchyan, Raveen Rathnasinghe, Zeyu Zhu, Simin Zheng, Nan Zhao, Kris White, Helen Ray-Jones, Valeriya Malysheva, Michiel J Thiecke, Siu-Ying Lau, Honglian Liu, Anna Junxia Zhang, Andrew Chak-Yiu Lee, Wen-Chun Liu, Teresa Aydillo, Betsaida Salom Melo, Ernesto Guccione, Robert Sebra, Elaine Shum, Jan Bakker, David A. Kaufman, Andre L. Moreira, Mariano Carossino, Udeni B R Balasuriya, Minji Byun, Emily R Miraldi, Randy A Albrecht, Michael Schotsaert, Adolfo Garcia-Sastre, Sumit K Chanda, Anand D Jeyasekharan, Benjamin R TenOever, Mikhail Spivakov, Matthew T Weirauch, Sven Heinz, Honglin Chen, Christopher Benner, Juergen A Richt, Ivan Marazzi

## Abstract

The ongoing pandemic caused by Severe Acute Respiratory Syndrome Coronavirus 2 (SARS-CoV-2) is currently affecting millions of lives worldwide. Large retrospective studies indicate that an elevated level of inflammatory cytokines and pro-inflammatory factors are associated with both increased disease severity and mortality. Here, using multidimensional epigenetic, transcriptional, *in vitro* and *in vivo* analyses, we report that Topoisomerase 1 (Top1) inhibition suppresses lethal inflammation induced by SARS-CoV-2. Therapeutic treatment with two doses of Topotecan (TPT), a FDA-approved Top1 inhibitor, suppresses infection-induced inflammation in hamsters. TPT treatment as late as four days post-infection reduces morbidity and rescues mortality in a transgenic mouse model. These results support the potential of Top1 inhibition as an effective host-directed therapy against severe SARS-CoV-2 infection. TPT and its derivatives are inexpensive clinical-grade inhibitors available in most countries. Clinical trials are needed to evaluate the efficacy of repurposing Top1 inhibitors for COVID-19 in humans.

## INTRODUCTION

Coronavirus disease 2019 (COVID-19) is an infectious disease caused by severe acute respiratory syndrome coronavirus 2 (SARS-CoV-2).

As of 16 November 2020, 54 million people have been infected and 1.33 million have died [https://coronavirus.jhu.edu/map.html, accessed 16 Nov 2020]. It is currently estimated that approximately 40-45% of SARS-CoV-2 infections are asymptomatic (Oran and Topol, 2020). For the remaining patients with symptomatic infections, data from China, Italy and the United States indicate that approximately 80% of infections are mild (not requiring hospitalization), 15% are moderate to severe (requiring hospitalization), and 5% are critical (requiring intensive care unit (ICU) care)(Garg et al., 2020; Livingston and Bucher, 2020; Stokes et al., 2020; Wu and McGoogan, 2020).

The most common manifestation of severe COVID-19 is acute hypoxemic respiratory failure, often associated with shock or multi-organ failure (Bhatraju et al., 2020; Wang et al., 2020a; Wu et al., 2020). Shock and multiorgan failure may be related to complications of critical illness; For example, ventilator-associated lung injury (Slutsky and Ranieri, 2013), secondary infection (Yang et al., 2020a) and aggravation of underlying chronic organ dysfunction(Cummings et al., 2020).

In most countries, the infection fatality rate is around 0.5-1%, and exponentially increases to up to 10% when referencing more susceptible age groups and people with pre-existing conditions(O’Driscoll et al., 2020). Therefore, the development of effective treatment plans for severe COVID-19 is imperative.

SARS-CoV-2 displays similar pathogenic mechanisms to that of SARS-CoV-1 (Channappanavar and Perlman, 2017; Zhu et al., 2020). The main hallmarks of disease progression feature two phases: a first phase of increasing viremia, followed by a subsequent steep increase in systemic inflammation (Lee et al., 2020; Merad and Martin, 2020; Siddiqi and Mehra, 2020). In fact, SARS-CoV-1 and SARS-Cov-2 patients who require intensive care showed elevated plasma levels of inflammatory cytokines and chemo-attractants (Chen et al., 2020; Del Valle et al., 2020; Huang et al., 2020; Lucas et al., 2020; Qin et al., 2020; Wang et al., 2020b; Yang et al., 2020b; Zhou et al., 2020a)

Several studies have shown that levels of inflammatory molecules can help distinguish those that survive COVID-19 from those that do not. For example, increased levels of IL-6, fibrin degradation products (D-dimer), as well as other single measurements like CRP or combined-measurement parameters (SOFA score) have been correlated with risk for death from COVID-19. (Zhou et al., 2020a). Notably, all non-survivors experienced sepsis (Zhou et al., 2020a). Therefore, increased systemic inflammation, occurring during disease progression, provides a biological rationale for interrupting hyper-inflammation to reduce disease severity. Guided by this logic, clinical trials have begun to examine the efficacy of cytokine blockers and anti-inflammatory molecules as potential COVID-19 therapeutics (Merad and Martin, 2020).

However, inhibition of single cytokines such as IL-6 or GM-CSF might not be sufficient (Hermine et al., 2020; Salvarani et al., 2020). This is due to the fact that many signaling molecules and pathways are involved in triggering an inflammatory response. Additionally, levels of individual cytokines can vary depending on the age and the clinical history of the patient, thus limiting the scope of therapeutics that only target a single inflammatory molecule.

A rapid induction of gene expression serves as a fundamental mechanism in activating an inflammation response. Inhibition of this process might hold the key to the development of novel therapeutics for COVID-19. Previously, we have reported that chromatin factors play key roles in controlling the induction of inflammatory gene expression programs (Marazzi et al., 2012; Miller et al., 2015; Nicodeme et al., 2010). Targeting the activity of these proteins acting on the chromatin template, where infection-induced gene transcription is executed, leads to the concerted suppression of multiple antiviral and anti-inflammatory genes (Marazzi et al., 2012; Miller et al., 2015; Nicodeme et al., 2010). Such simultaneous inhibition of many virus-induced genes “in one go” can have a clear advantage over conventional single target therapies (Marazzi et al., 2018). In particular, we have previously shown that the host enzyme topoisomerase 1 (Top-1) is required to selectively activate the expression of inflammatory genes during viral and bacterial infection and co-infections (Rialdi et al., 2016). Therapeutic administration (after infection) of one to three doses of topoisomerase inhibitors can rescue mortality in four animal models of inflammation-induced death (Rialdi et al., 2016). These data support the hypothesis that host-directed epigenetic therapy can be valuable to suppress hyper-inflammatory responses in the context of infectious diseases. We present here, a series of experiments in which we tested the hypothesis that epigenetic therapy aimed at modifying the host response to SARS-CoV-2 infection would ameliorate severe COVID-19.

## RESULTS

Cell signaling cascades converge on chromatin to dictate changes in gene expression programs upon cell intrinsic and extrinsic signals. We performed a combined structural and epigenetic analysis during infection in an effort to understand how SARS-CoV-2 alters chromatin function and the epigenetic landscape of cells upon infection,

To characterize structural chromatin changes we performed Hi-C, a technique that profiles the three-dimensional architecture of the genome (Bonev and Cavalli, 2016; Hildebrand and Dekker, 2020). Hi-C was performed on uninfected and SARS-CoV-2 infected A549 cells expressing the human SARS-CoV-2 entry receptor Ace2 (A549-ACE2) at both early (8 hours) and late (24 hours) time points post infection, in order to investigate the dynamics of infection induced changes on chromatin structure. Reproducible results were achieved across replicates for all time points. (**Table S1**). Our analysis indicates that large portions of the genome alter their global interaction profiles as infection progresses, culminating in a major redistribution of chromatin associated with either the active (A) or inactive (B) compartments at the 24h time point (**Figure 1A**). Notably, compartment changes result in a shortening of the domain size, with large linear stretches of A and B compartment chromatin generally becoming subdivided into smaller A/B domains (**Figure 1A and 1B**), a feature that partially phenocopies the loss of regional topological constraints controlled by cohesin (Rao et al., 2017; Schwarzer et al., 2017).

**Figure 1:**
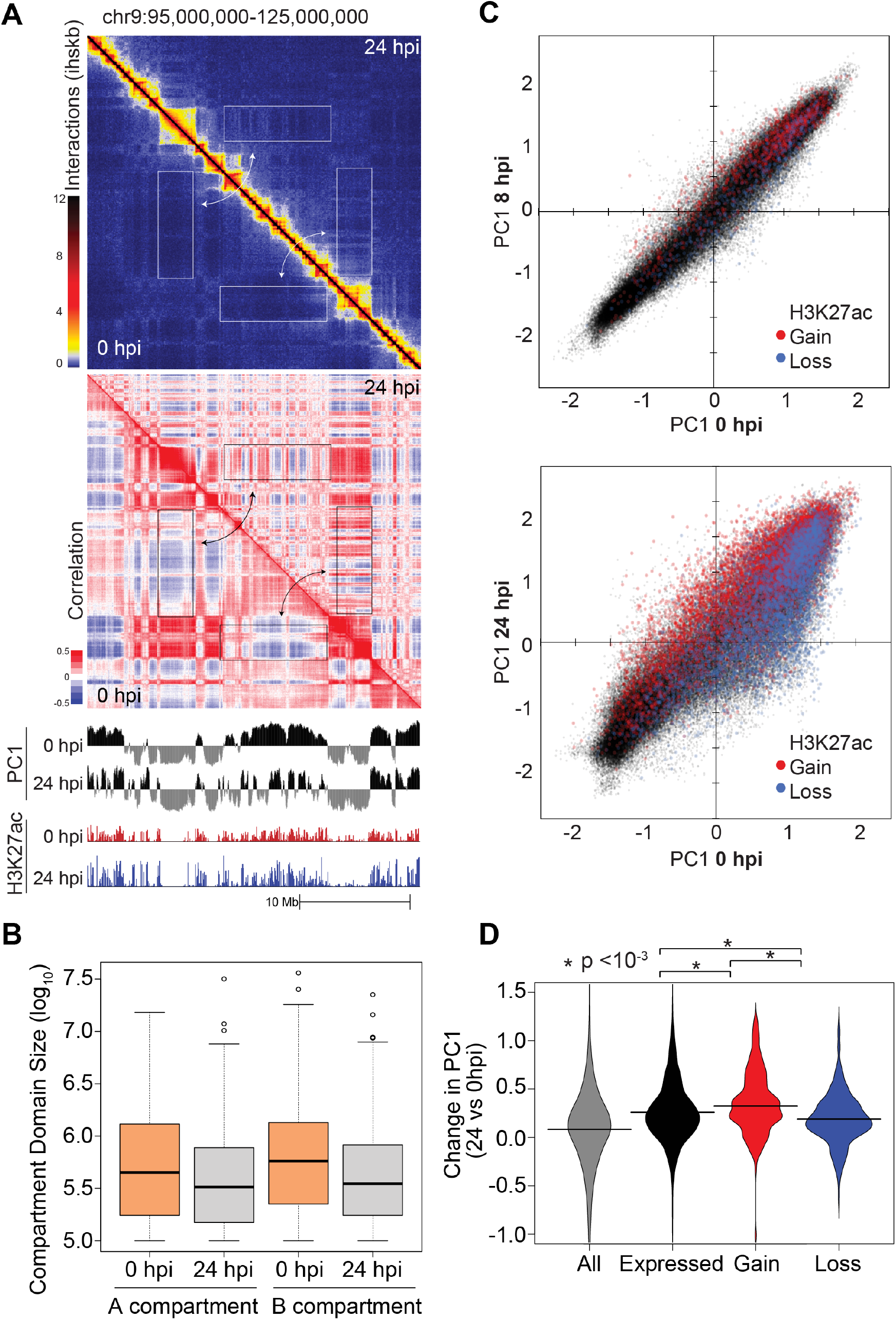
SARS-CoV-2 restructures chromatin in host cells. **(A) Upper panel**: Analysis of 3D chromatin structure in SARS-CoV-2 infected and control A549-ACE2 cells by Hi-C. Normalized Hi-C contact matrices are shown for the uninfected (0hpi) control (lower-left) and 24 hours post infection (hpi, upper-right) for a representative 30 Mb region of chromosome 9. White rectangles highlight regions with strong changes in interaction patterns between conditions. **Middle panel**: pairwise correlation matrices for uninfected control and 24 hpi Hi-C experiments analysis for the same region shown in the upper panel. **Lower panel**: PC1 values, which represent the PCA loadings describing the chromatin compartment membership (+ values for the A compartment, - values for the B compartment) are show along with H3K27ac ChIP-seq levels for the region depicted. Cells infected for 24 hours show increased segregation of chromatin into smaller A and B compartment domains. **(B)** Distribution of A and B compartment domain sizes genome wide for uninfected control and 24 hpi A549-ACE2 cells. **(C)** Scatter plot comparing the PC1 values for every 25 kb region in the genome for uninfected control and infected cells (8, 24 hpi). Data points colored red or blue indicate that they overlap with a significantly regulated H3K27ac peaks (4-fold, adjusted p-value < 0.05). **(D)** Distribution of the change in PC1 values between uninfected and 24 hpi at the promoters of genes that are either expressed in A549-Ace2 cells, induced, or repressed by SARS-CoV-2 infection (>1.5-fold, adjusted p-value < 0.05).

To characterize whether the structural changes seen using Hi-C were associated with epigenetic features of gene activation and repression, we performed ChIP-sequencing for H3K27 acetylation (K27ac), an epigenetic mark found at active regulatory regions that is commonly used to monitor dynamic changes in transcriptional activation. ChIP-seq for H3K27ac was performed in uninfected and infected A549-ACE2 cells at the same time points as the Hi-C. Our analysis showed high correlation of K27ac levels between replicates (**Figure S1A**). While some regions of the genome showed no change in K27ac levels upon infection (cluster i; **Figure S1B and S1C**), there were statistically significant changes in K27ac levels at promoters and other regulatory regions during the course of infection (clusters ii to vii; **Figure S1B and S1C; Table S2A-S2C**). Regions that significantly gain (clusters v and vi; **Figure S1B**) and lose K27ac (clusters ii and iii; **Figure S1B**) over the course of infection were detected.

We labelled changes in K27ac across condition as “K27 loss” to describe instances in which reduction of K27ac signal across the genome occurs during infection compared to the uninfected condition. Increased genomic level of K27ac during infection was labelled as “K27 gain”.

Overlaying the changes in K27ac with structural chromatin changes indicate that regions gaining or losing K27ac are enriched in chromatin domains that move from B-A or A-B compartment, respectively (**Figure 1C**). This partitioning occurs dynamically throughout the infection (**Figure 1C**-compare 8h vs 24h) and is associated with gene expression activity (**Figure 1D**). These results suggest that the dynamic restructuring of genome compartmentalization by SARS-CoV-2 infection is highly associated with transcriptional activity.

To characterize whether a unique set of transcription factors might participate in partitioning chromatin into active or inactive regions during infection, we performed DNA binding site motif enrichment analysis of regions displaying differential H3K27ac activity. Our results indicate that repressed regions lack unique enrichment of immune-specific transcription factors at promoters, enhancers, and other putative regulatory regions (**Figure S1D, Table S3**). Regions that gain K27ac signal are largely devoid of unique transcription factor signatures apart from a strong enrichment for motifs recognized by NF-kB (red bars, cluster v and vi; **Figure S1D, Table S3**), a master regulator of inflammatory gene programs.

Overall, our results indicate that SARS-CoV-2 induces global epigenetic and structural changes in the infected cell, underlying the identity of inducible regulatory networks controlling cell responses to infection.

### Topoisomerase I controls SARS-CoV-2-induced gene expression response

To determine whether chromatin factors can control the induction of the cell response by promoting the transactivation of SARS-CoV-2-induced gene program, we focused our attention on topoisomerase I (Top1), a factor known to activate bacterial- and viral infection-induced genes (Rialdi et al., 2016). We performed siRNA-mediated knockdown of Top1 (siTOP1) along with control siRNA (siSCR) in A549-ACE2 cells, followed by mock treatment (PBS only; uninfected controls) or infection with SARS-CoV-2. Gene expression changes in these cells were quantified by RNA-sequencing at 24 hours post infection. Our analyses indicate that siTOP1 treated cells had a distinct transcriptional response to the virus (**Figure 2A**) as compared to no siRNA or siSCR treated cells, and that depletion of Top1 resulted in selective suppression of many infection-induced genes (Fold Change > 1.5, padj<0.05; **Figure 2B and 2C, Table S4**). Gene ontology pathway analyses of genes that are downregulated upon Top1 knockdown suggest that many of these genes are involved in inflammatory responses (**Figure 2C, Table S5**). We further validated our results by qPCR for representative genes IL-6, CXCL8 and TOP1 (**Figure 2D**), verifying that depletion of Top1 reduces the expression of these inflammatory genes.

**Figure 2:**
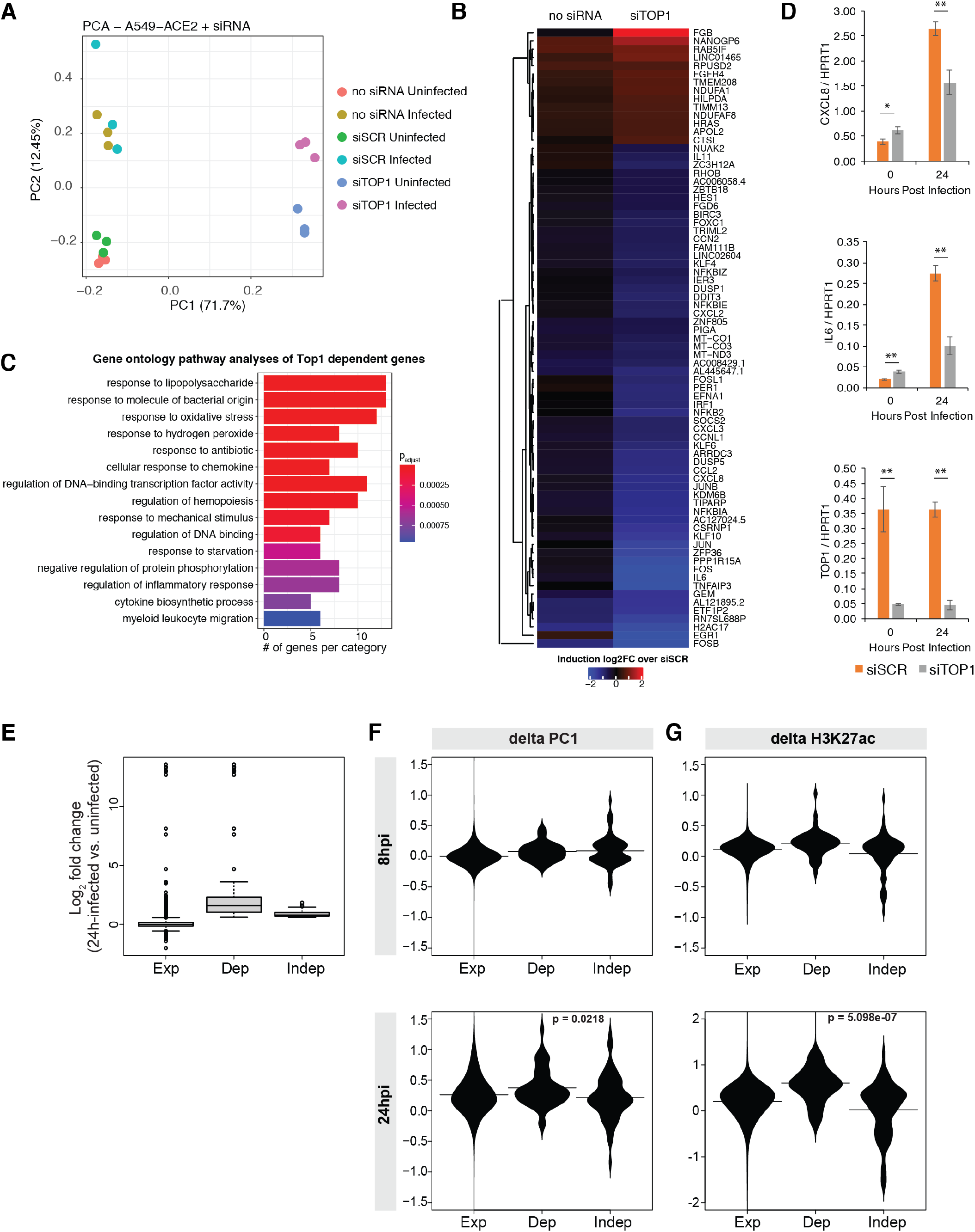
TOP1 depletion in SARS-CoV-2 infected cells inhibits induction of inflammatory genes. **(A)** PCA plot showing the relationship between samples and treatment conditions. **(B)** Heat map showing relative changes in gene expression levels in no siRNA (no siRNA) or siTop1 (siTOP1) treated cells, when compared to nontargeting control siRNA-treated (siSCR) cells. Shown are genes that are differentially expressed between siTOP1 and siSCR samples (adjusted p-value < 0.05, fold change>1.5). **(C)** Gene ontology analyses of downregulated target genes shown in (B). **(D)** qPCR validation of select target genes shown in (B). Shown are the mean and s.d of 3 replicates. *: p<0.05; **: p<0.01 by two-tailed, unpaired Students’ t-test. **(E)** Barplots showing changes in gene expression levels upon SARS-CoV-2 infection, as quantified by RNA seq, for all expressed genes (Exp), Top1 dependent induced genes (Dep) and Top1-independent induced genes (Indep) **(F)** Violin plots showing changes in PC1 (delta PC1) for 8 hours (8hpi) and 24 hours (24hpi) post infection at expressed genes (Exp), Top1 dependent induced genes (Dep) and Top1-independent induced genes (Indep). Horizontal lines indicate the means. **(G)** Violin plots showing changes in H3K27ac levels (delta H3K27ac) for 8 hours (8hpi) and 24 hours (24hpi) post infection at expressed genes (Exp), Top1 dependent induced genes (Dep) and Top1-independent induced genes (Indep). Horizontal lines indicate the means.

To understand the specificity of Top1, we profiled infection induced and Top1 dependent genes (“Dep.”; **Table S4C**) identified in Figure 2B with respect to their structural and epigenetic status at basal state and after infection. As controls, we used all expressed genes (“Exp”; **Table S4C**) or genes that are also induced by infection but unaffected by Top1 depletion (Top1 independent, “Indep.”; **Table S4C**). Our analysis indicates that genes that depend on Top1 for their upregulation are induced to higher levels then Top1 independent genes upon infection (**Figure 2E**). Top1 dependent genes also displayed greater shifts towards active chromatin compartment (positive delta PC1 levels, **Figure 2F**) and increases in K27ac signals (positive delta H3K27ac levels; **Figure 2G**) compared to Top1 independent genes. Differences were more pronounced later in infection (**Figures 2F and 2G**).

Overall our analysis indicates that Top1-dependent genes are highly inducible during SARS-CoV-2 infection as a result of chromatin and epigenetic changes that render their transactivation permissible.

### Top1 inhibition suppresses lung inflammation and lung damage in infected hamsters

To determine whether inhibition of Top1 activity can dampen inflammatory gene expression *in vivo*, we selected topotecan (TPT), a FDA approved Top1 inhibitor, to use in the Syrian Golden hamster model (Munoz-Fontela et al., 2020) (hereafter referred as hamster) of SARS-CoV-2 infection.

We treated SARS-CoV-2 infected hamsters with either vehicle control (DMSO) or 10mg/kg TPT at days 1 and 2 post-infection. Lungs from these animals were then collected for histology and transcriptome analysis at days 4 and 6 post-infection (**Figure 3A**).

**Figure 3:**
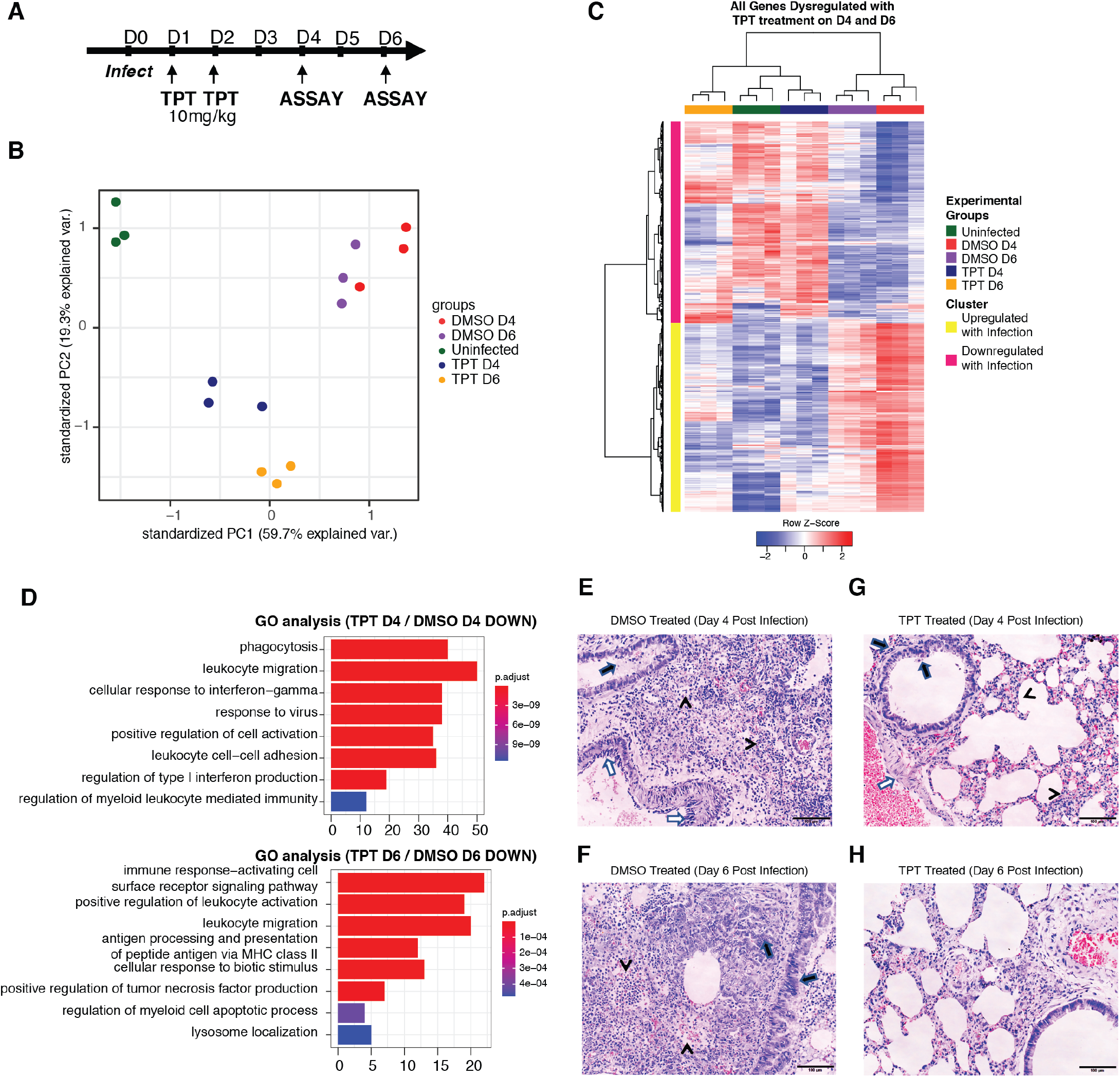
TPT treatment reduces inflammatory gene expression in SARS-CoV-2 infected hamsters. **(A)** Schematic showing the infection and treatment regime used. **(B)** PCA plot showing the relationship between treatment and infection groups. **(C)** Heatmap showing gene expression levels of genes that are dysregulated with TPT treatment in Uninfected (Green), DMSO (red and purple) or TPT treated (blue and yellow) hamsters at days 4 and 6 post infection. Each column represents an individual hamster, and each row represents one gene. Data are clustered by genes that are up- (yellow) or down- (pink) regulated with infection with reference to the uninfected animals. **(D)** Gene ontology analysis of genes that are down regulated with TPT treatment at days 4 (top) and 6 (bottom) post infection **(E)** Representative scan of hematoxylin and eosin (H&E) stained sections of the lungs of infected hamsters that have been treated with DMSO 4 days post infection. Arrow: Diffuse lung inflammatory damages. Bronchiolar epithelium cells death, bronchiolar luminal secretion and hemorrhage; Arrowheads: Diffuse alveoli destruction with massive immune cell infiltration and exudation; Open arrows: Vasculitis **(F)** Representative scan of hematoxylin and eosin (H&E) stained sections of the lungs of infected hamsters that have been treated with DMSO 6 days post infection. Lung tissue consolidation affected most of the lung lobe examined. Bronchial secretion, infiltration and alveolar space exudation, immune cell infiltration and hemorrhage are still present at this stage (arrowheads), and is accompanied by alveolar and bronchiolar cell proliferation (arrows) **(G)** Representative scan of hematoxylin and eosin (H&E) stained sections of the lungs of infected hamsters that have been treated with TPT 4 days post infection. Diffuse milder inflammatory damages. Arrows: Bronchiolar epithelium cells death with milder peribrochiolar infiltration; Arrowheads: Diffuse alveolar wall thickening with capillary congestion. No conspicuous alveolar space infiltration, exudation or hemorrhages; Open arrows: Vasculitis is very mild and rare **(H)** Representative scan of hematoxylin and eosin (H&E) stained sections of the lungs of infected hamsters that have been treated with TPT 6 days post infection. Patchy lung tissue consolidation with cell proliferation. Most alveolar area are without exudation and infiltration. Bronchiolar luminal secretion is reduced compared to the with DMSO control.

Clustering of RNA-seq reads using principal component analysis (PCA) indicates that the gene expression profiles under the three conditions (uninfected, infected-DMSO treated, and infected-TPT treated) partition based on infection, treatment status, and the temporality of the infection (day 4 and 6), with each replicate clustering in close proximity to its counterpart (**Figure 3B**).

Differential expression (DE) analysis showed that TPT suppresses inflammatory gene expression in the lungs of infected hamsters (**Figure 3C and 3D**). Clustering of the DE data indicates that the gene expression profiles of TPT-treated infected lungs are more similar to that of the non-infected lungs, rather than infected ones (**Figure 3C**). The GO categories associated with the TPT-suppressed genes indicates specific inhibition of virus-induced and inflammatory genes at both day 4 and day 6 post-infection (**Figure 3D**).

Histopathological analysis of infected, DMSO vehicle-treated hamster lungs at days 4 and 6 post-infection displayed diffused alveoli destruction, bronchiolar epithelium cell death and hemorrhaging, coupled with massive immune cell infiltration and exudation, typically associated with increased expression of inflammatory mediators and recruitment of immune cells during infection (**Figure 3E and 3F**). On the contrary, TPT treatment diminished pathological features of lung damage in infected animals. Lungs from these animals did not have conspicuous alveolar space infiltration, exudation or hemorrhaging at both days 4 and 6 post infection (**Figure 3G and 3H**).

To determine the clinical significance of our observations, we then asked if the genes that were downregulated by TPT treatment in SARS-CoV-2 infected hamsters also corresponded to immunopathological gene signatures that have been observed in COVID-19 patients. Cross-comparison of our results with the gene expression profiles in human lungs isolated from autopsies of COVID-19 patients and uninfected control lungs (Nienhold et al., 2020) indicated that TPT suppressed genes that are hyperactivated in patients who succumbed to infection (**Figure 4A and 4B**). In fact, TPT-inhibited genes are up-regulated in COVID-19 lung autopsy tissue relative to healthy control (P<1E-3) (**Figure 4A, left panel**), while genes up-regulated by TPT are down-regulated in COVID-19 lung relative to control (P<1E-7) (**Figure 4A, right panel**). These results suggest that treatment with TPT might reverse COVID-19-induced lung gene expression responses. The gene expression profiles of TPT inhibited genes in individual patients (**heatmap, Figure 4B**) and the corresponding gene set enrichment scores are shown in **Figure 4B**.

**Figure 4:**
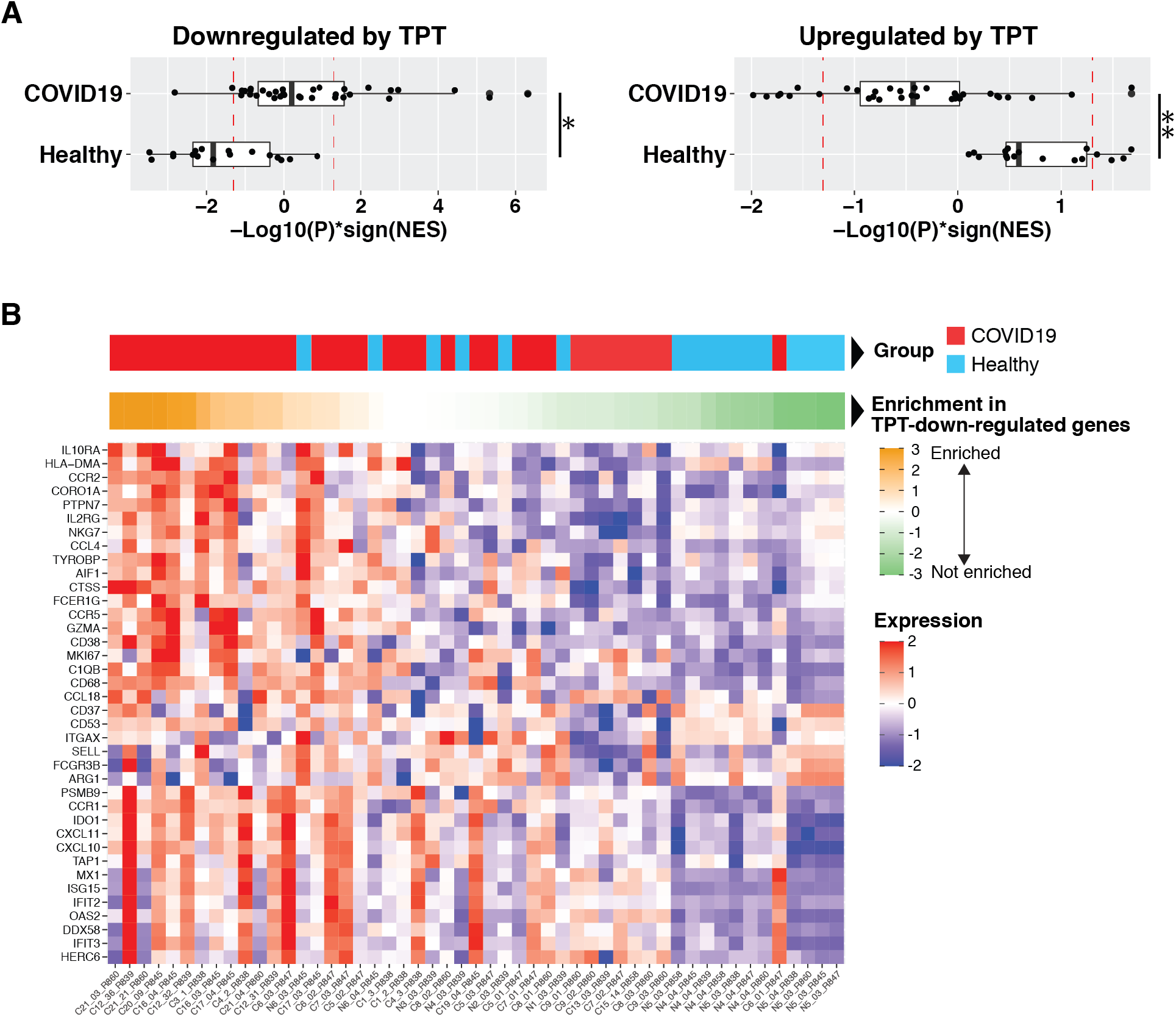
TPT suppresses gene programs upregulated in autopsy-lung from COVID19 patients. **(A)** Gene set enrichment analysis of lung-tissue gene expression profiles from COVID19 deceased patients versus healthy patients (Nienhold et al., 2020). Signed –log10 adjusted P values indicate enrichment of down-regulated (top panel) and up-regulated (bottom panel) gene signatures from TPT-treated hamsters infected with SARS-CoV-2. The sign of enrichment is given by the normalized enrichment score (NES). Dashed lines indicate a significance levels of P=0.05. Differences in mean NES are shown: * P = 10-3, ** P = 5×10-4, *** P = 10-7. **(B)** Expression in lung autopsy tissue of COVID19 patients and healthy controls [(Nienhold et al., 2020), GSE151764] of genes down-regulated in TPT-treated Sars-CoV-2-infected hamsters (log2|FC| > 1, FDR=10%). Patient groups are indicated by the topmost bar, where healthy controls are colored in cyan, and COVID-19 patients colored in red. Gene set enrichment scores, calculated as −log10(P)*sign(NES) are indicated in the middle bar. The sign of enrichment is given by the normalized enrichment score (NES). Positive, higher scores (orange) indicate that TPT-inhibited genes are more upregulated in a given patient, whereas negative, lower scores (green) indicate that TPT-inhibited genes are more downregulated in a given patient. The lower heatmap shows the individual gene expression profile of the indicated TPT-inhibited gene for a given patient (in columns). Heatmap is sorted by column from the highest (left) to lowest enrichment score (right).

We next sought to validate whether lower dosages of TPT, which are associated with negligible cytostatic effects (Guichard et al., 2001; Houghton et al., 1995; Nemati et al., 2010), were effective in suppressing SARS-CoV-2 infection induced inflammation. We performed a parallel experiment to the one described in **Figure 4A** using 5-fold lower TPT (2mg/kg) and the same regimen of TPT treatment at Day 1 and 2 post-infection (**Figure S2A**). Lungs from infected and treated hamsters were assayed at Day 4 post infection.

Animals treated with TPT had reduced lung to body weight ratios post infection (**Figure S2B**), which suggest reduced pulmonary edemas in these animals. In line with this, histopathological analyses showed reduced broncho-pneumonia (**Figure S2C-S2E**) and immune cell infiltration (**Table S6**) in the lungs of TPT treated animals when compared to DMSO treated ones. qPCR analysis of representative genes also suggested reduced expression of inflammatory genes in TPT treated animals (**Figure S2F**). Overall, these results suggest that lower doses of TPT treatment can still effectively suppress the expression of inflammatory molecules, and ameliorate inflammation induced pathology during SARS-CoV-2 infection.

In sum, our results support the hypothesis that TPT suppresses SARS-CoV-2-induced lung inflammation in vivo.

### Top1 inhibition therapy suppresses, SARS-CoV-2 morbidity and lethality in transgenic mice

To further verify our results, we extended our studies to a complementary model and evaluated the effects of TPT treatment in transgenic mice that express the human angiotensin I-converting enzyme 2 (ACE2) receptor under the cytokeratin 18 gene promoter (K18-hACE2). This mouse strain is susceptible to SARS-CoV-2 infection and displays a disease progression profile that shares many features of severe COVID-19 (Winkler et al., 2020). Importantly, loss of pulmonary function and weight loss in these mice occurs after the peak of viral replication, and coincides with infiltration of immune cells (monocytes, T cells, neutrophils) in the lung and alveolar spaces at day 4 post infection (Winkler et al., 2020). As such, K18-hACE2 have been suggested as a model to define the basis of SARS-CoV-2-induced lung disease and test immune and antiviral countermeasures (Bao et al., 2020; Winkler et al., 2020).

To test whether inhibition of inflammation provides a protective effect in infected K18-hACE2 mice, we performed three different regimes of TPT treatments, labelled as early, intermediate, and late, to respectively describe dosing of the inhibitor at 2mg/kg on days 1+2, days 3+4; or days 4+5 post-infection respectively (**Figure 5A**).

**Figure 5:**
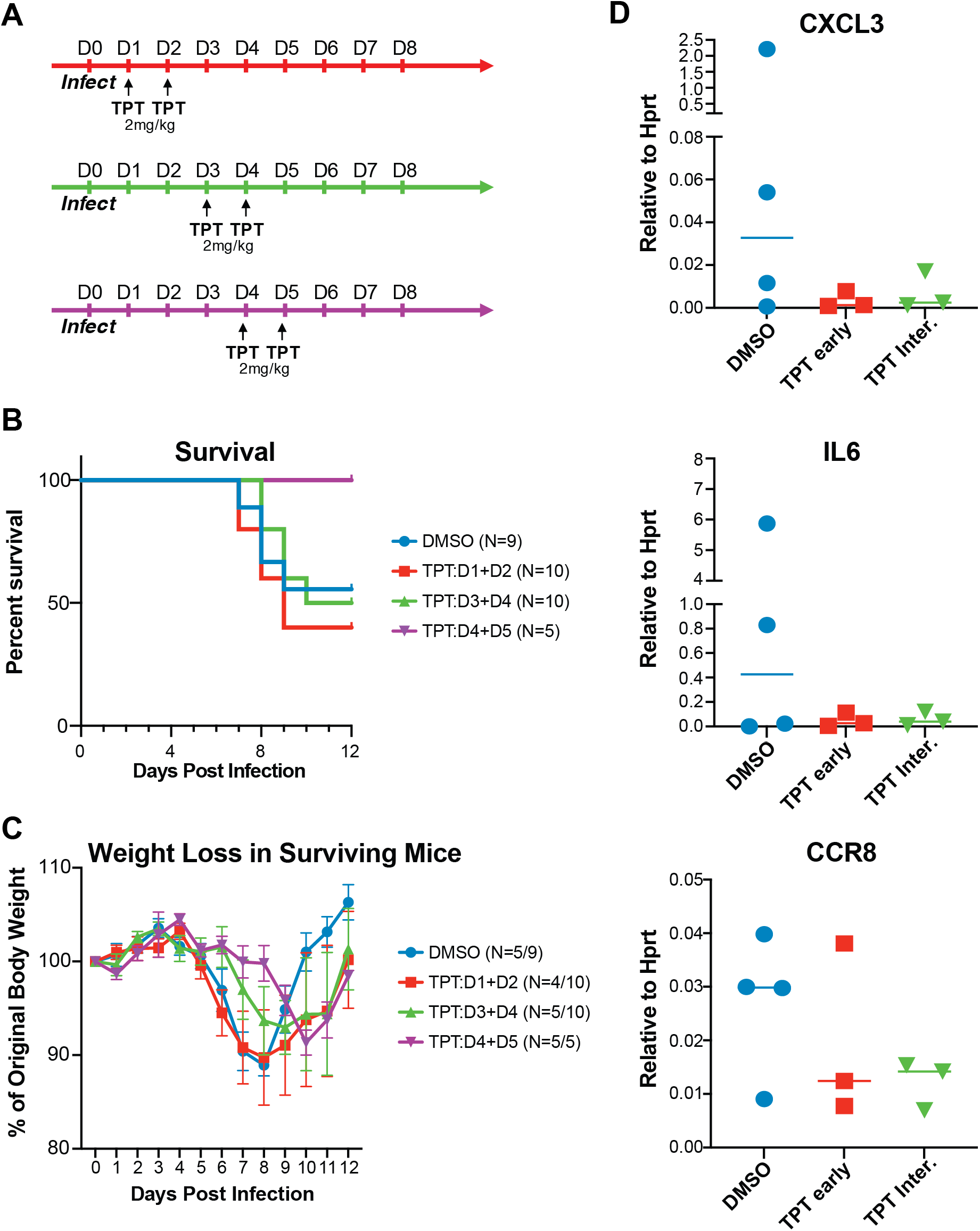
Late treatment of TPT in K18-hACE2 mice provides survival benefit during SARS-CoV-2 infection. **(A)** Schematic showing infection and treatment regime in mice. Groups are color coded by treatment regime **(B)** Survival curve of K18-hACE2 mice infected with 1E4 PFU of SARS-CoV-2 and subsequently subjected to the indicated TPT treatment regimes. Number of mice used are indicated in the legend. Blue: DMSO vehicle control only (n=9); Red: TPT 2mg/kg on Days 1 and 2 post infection (n=10); Green: TPT 2mg/kg on Days 3 and 4 post infection (n=10); Purple: TPT 2mg/kg on Days 4 and 5 post infection (n=5). **(C)** Weight loss curves in surviving mice shown in B. Numbers of mice at the end and start (end/start) points of the experiment are indicated in the legend keys. Weights are shown as means of the percentage of starting weights. Error bars show the SEM of each group. Blue: DMSO only; Red: TPT 2mg/kg on Days 1 and 2 post infection; Green: TPT 2mg/kg on Days 3 and 4 post infection; Purple: TPT 2mg/kg on Days 4 and 5 post infection. **(D)** qPCR of inflammatory gene expression for mice in the indicated treatment groups at day 7 post infection with 1E4 PFU SARS-CoV-2. Each dot represents an individual mouse. Lines indicate the mean of expression.

The rationale behind this approach is that inhibition of inflammation could be detrimental during the early phases of the infection, as it might cause increased viral replication and dissemination. The optimal protective effect of inhibiting inflammation should be achieved during the hyper-inflammatory phase of the disease, which would coincide with the later stage of infection.

Indeed, our results showed that early treatment of TPT is ineffective in reducing the morbidity and mortality caused by SARS-2 infection (**Figure 5B and 5C: TPT: D1+D2**). Intermediate treatment ameliorates morbidity but not mortality (**Figure 5B and 5C: TPT: D3+D4**). Strikingly, late TPT treatment suppresses both morbidity and mortality (**Figure 5B and 5C: TPT: D4+D5**). TPT treatment is associated with suppression of inflammatory gene expression in the lung, as indicated by qPCR of representative genes IL-6, CXCL3 and CCR8 (**Figure 5D**).

Our results indicate that inhibition of hyper-inflammation by therapeutic administration of TPT can rescue K18-hACE2 mice from lethal SARS-CoV-2 infection.

## DISCUSSION

The ongoing COVID-19 pandemic caused by SARS-CoV-2 is currently affecting millions of lives worldwide, and poses an overwhelming burden on global health systems as well as the economy. The development of novel therapeutics against SARS-CoV-2 remains a top priority. While prophylactic measures are being evaluated and distributed, drugs available to target SARS-CoV-2 and function therapeutically are direly needed especially for severe cases of COVID-19.

Although the pathophysiology of SARS-CoV-2 has not yet been fully characterized, it has been observed that SARS-CoV-2 infection triggers hyper-activation of pro-inflammatory cytokines (IL-6, -Il1b, TNFa) and chemokines (CXCL8, 9, 10, CCL2) (Huang et al., 2020; Lucas et al., 2020; Merad and Martin, 2020; Tang et al., 2020; Zhou et al., 2020a). The increased level of inflammatory molecules has been shown to correlate with COVID-19 disease severity (Del Valle et al., 2020; Moore and June, 2020). While the exact mechanism and cell type-specific contributions to hyper-inflammation still needs to be fully elucidated, monocytes, macrophages and dendritic cells are primary candidates and have been reported to contribute to the cytokine-mediated immunopathology seen in human (Del Valle et al., 2020; Giamarellos-Bourboulis et al., 2020; Moore and June, 2020). This is supported by previous studies of the immune response against SARS-CoV-1 and MERS-CoV infections (Cheung et al., 2005; Wong et al., 2004). Additionally, non-myeloid cells have been recently shown to contribute to the hyper-inflammatory program (Zhou et al., 2020b). Elevated inflammatory response contributes to sepsis and multi-organ failure, an important contributor to death from COVID-19 (Zhou et al., 2020a). Therefore, treatments that can suppress host inflammatory response might be potentially effective therapeutic strategies for COVID-19. In this light, it is important to highlight that glucocorticosteroids (dexamethasone, methylprednisolone, hydrocortisone), which act as suppressors of systemic inflammation, have been reported to ameliorate the outcome of COVID-19, especially in hospitalized patients who require supplemental oxygen (Group et al., 2020).

### Structural and functional changes in the genome upon infection

While our knowledge of SARS-CoV-2 pathogenesis is expanding rapidly, little is known about how epigenetic modifications and genome structure are affected by infection, and in what capacity they affect gene activity (Liu et al., 2020). Our data suggest that SARS-CoV-2 infection imposes a pervasive effect on the host cell response at the genome organization level, causing drastic changes in both the segmentation and interaction patterns of chromatin regions. The SARS-CoV-2 infection-induced chromatin segregation pattern mirrors what observed in cells depleted of cohesin, a protein that maintains chromosomal architecture (Hnisz et al., 2016; Schwarzer et al., 2017). In both SARS-CoV-2 infection and absence of cohesin, large stretches of A and B compartments become subdivided into smaller A/B domains, leading to reduction in compartment domain size. These effects are likely indicative of increased chromatin fiber flexibility, allowing it to segregate more easily according to the activity of the embedded regulatory elements. Future work is needed to understand the implications of this events for infection response.

Regarding SARS-CoV-2 infection-induced compartment movement, we surmise that A-to-B and B-to-A switches are driven by transcriptional and epigenome activity. While A-B transitions are characterized by decreased K27ac at promoters and gene suppression, B-A is accompanied by increased K27ac at promoters and gene induction. Gene suppression has functional consequences as it affects many conventional infection-induced genes activated by STAT1/2 and IRF3 transcription factors. Suppression is likely a result of viral antagonism. In fact, a recent report elucidates the pleiotropy of viral antagonism mechanisms and their collective effects on suppressing host functions at transcriptional, co-transcriptional and post-transcriptional levels (Banerjee et al., 2020; Lei et al., 2020). Gene activation is the result of signal-induced transactivation, and indicates that many cellular genes escape viral suppression during infection. One prominent example is a subset of inflammatory genes whose expression is driven by infection-activated transcription factor NF-kB. The proteins encoded by these genes are potent pro-inflammatory molecules and present systemically with high levels in severe COVID-19 patients (Del Valle et al., 2020; Moore and June, 2020). The selective and concerted induction of inflammatory genes provides the rationale for using epigenetic inhibitors to suppress their induction and establish a global anti-inflammatory state (Marazzi et al., 2018).

### Top1 inhibition therapy

We show that the host enzyme Topoisomerase-1 promotes transcriptional activation of pro-inflammatory genes during SARS-CoV-2 infection. We then demonstrate that Top1 inhibition limits the expression of inflammatory genes in the lungs of infected animals. Most importantly, Top1 inhibition decreases morbidity and morbidity in infected mice. The therapeutic effect can be achieved by drug administration 4-5 days following infection.

Whether the suppression of inflammation *in vivo* is solely the result of dampened epithelial response, or whether it affects, directly or indirectly neutrophil/monocytic activation or immune cell recruitment to the lung remains unknown. We posit that TPT action in both epithelial and immune cells responses result in the positive outcomes, as TPT is likely to suppress inducible transcriptional programs in both the infected cells and by-stander cells. Dampening highly inducible genes and sparing housekeeper genes is a typical feature of epigenetic inhibitors that act on signal-induced genes, which aside from the requirement of cofactors for their activation have unifying genomic features like high burst rates conferred by many regulatory enhancers(Chen et al., 2019; Fukaya et al., 2016; Marazzi et al., 2018; Senecal et al., 2014; Zabidi et al., 2015).

In sum, TPT and other Top1 inhibitors like irinotecan are widely available and FDA-approved. Some are in the WHO list of Essential Medicines. They are inexpensive and generic formulation exists throughout the world, making them easily accessible for immediate use. Overall, our results suggest that repurposing of TOP1 inhibitor could be a valuable strategy to ameliorate or treat severe COVID-19.

### Pharmacokinetics considerations and limitations of this study

Although the preclinical animal models of SARS-CoV-2 pathogenesis used here are, as with any animal model, only partially representative of the biology of humans, our study indicates a promising effect of TPT by suppressing inflammation in COVID-19. Several factors require careful consideration prior to extrapolating these results towards the design of clinical trials of Top1-inhibition therapy for human COVID-19. First, in our animal models, we can suppress inflammation and reduce disease pathology in the lung using 2 doses of Top1 inhibition therapy with TPT at 2mg/kg intraperitoneally. This equates to a 5-fold reduction from typical chemotherapeutic anti-cancer doses in rodent models (Guichard et al., 2001; Houghton et al., 1995; Nemati et al., 2010). In clinical practice, the Top1 inhibitors TPT and Irinotecan have well-characterized pharmacokinetics and toxicity profiles (Kollmannsberger et al., 1999; Mathijssen et al., 2001), albeit in patients without SARS-CoV-2 infection. Doses that are 5-fold lower than those used in the treatment of small-cell lung cancer (TPT)(Rowinsky et al., 1992; von Pawel et al., 1999) and colorectal cancer (irinotecan)(Andre et al., 1999) are expected to cause little to no toxicity, and importantly no risk of neutropenia. This significant dose reduction, together with the wealth of clinical experience in the use of TPT and irinotecan should reassure us about potential concerns over cytotoxicity. Nonetheless, safety trials of the reduced dosage of TPT or irinotecan in COVID-19 patients will need to be performed prior to testing efficacy. Another important consideration is that the window of opportunity for Top1 inhibitor treatment in humans needs to be carefully evaluated. Many reports indicate that the timing of the intervention against Coronaviruses is key, as protective anti-viral and damaging excessive inflammatory responses need to be balanced (Channappanavar et al., 2016; Channappanavar et al., 2019; Grajales-Reyes and Colonna, 2020). Our data aligns with those studies, as early treatment of TPT did not display protective effect in mice. In essence, limiting inflammatory response too early during infection might increase viral replication and dissemination. As TPT-mediated inhibition of inflammation could theoretically lead to a resurge in viral replication, clinical trials will ideally need to incorporate the administration of an anti-viral agent with activity against SARS-CoV-2 after TPT treatment. The safety and efficacy of our strategy will now be evaluated in two clinical trials of TPT that have been submitted for trial initiation, and are set to begin in January 2021 (A.J, D.K., and I.M. -*personal communication*). Lastly, we strongly discourage any ‘off label’ use of Top-1 inhibitors until safety and effectiveness is established by clinical trials.

## AUTHOR CONTRIBUTIONS

Conceptualization: I.Marazzi

Investigation: J.S.Y.H., B.W.-Y.M., L.C., T.J., S.Y., S.P., J.A.W., N.N.G., D.A.M., S.V.I., I.Morozov, J.D.T., Y.S.F., R.R., Z.Z., S.Z., N.Z., B.S.M, H.R.-.J., V.M., M.J.T., S.-Y.L., H.L., A.J.Z., A.C.-Y.L., W.-C.L., T.A.-G., A.M., R.A.A., M.Schotsaert, S.H.

Genomics analyses (Epigenetics): S.P., J.A.W., E.R.M., M.T.W. Genomics analyses (Chromatin structure): S.H., C.B.

GSEA analysis: J.A.W., E.R.M.

Data analyses (Others): J.S.Y.H., Y.S.F., H.R.-J., V.M., M.Spivakov

In vivo study and veterinarian analysis: J.S.Y.H, B.W.-Y.M., L.C, S.-Y. L., H.L., A.J.Z., A.C.-Y.L., H.C., N.N.G., D.A.M., S.V.I., I.Morozov, J.D.T., J.A.R., M.C., U.B.R.B.

Histology and medical consultation: E.S., D.K., A.M., J.B., A.D.J.

Writing-Original draft: I.Marazzi

Resources: K.W., M.J.T., R.S., A.G.-S., B.R.T., S.K.C.

Writing-Review & Editing: J.S.Y.H, Z.Z., E.G., E.S., D.A.K., M.B., E.R.M., A.G.-S., M.T.W., S.H., C.B., J.A.R., I. Marazzi

Funding acquisition: A.G.-S., H.C., C.B., J.A.R., I. Marazzi

Project administration: I. Marazzi

Supervision: I. Marazzi

## ACKNOWLEDGEMENTS

We thank the staff of KSU Biosecurity Research Institute, the histological laboratory at the Kansas State Veterinary Diagnostic Laboratory (KSVDL), members of the Histology and Immunohistochemistry sections at the Louisiana Animal Disease Diagnostic Laboratory (LADDL), the CMG staff and Bianca Artiaga, Dashzeveg Bold, Konner Cool, Emily Gilbert-Esparza, Chester McDowell and Yonghai Li. We thank the teams from the Genomics and Mouse facilities at Icahn School of Medicine at Mount Sinai, the Global Health and Emerging Pathogens Institute (GHEPI) at Mount Sinai. We thank Alan Soto from the Biorepository and Pathology Dean’s CoRE at the Icahn School of Medicine at Mount Sinai. We thank Cindy Beharry, Sonia Jangra, Nanyi Julia Zhao, Nancy Francoeur, Nataly Fishman, Marion Dejosez, Thomas Zwaka and Carles Martinez-Romero for their help and advice.

This work was partially supported through grants from NBAF Transition Funds, the NIAID Centers of Excellence for Influenza Research and Surveillance under contract number HHSN 272201400006C and the Department of Homeland Security Center of Excellence for Emerging and Zoonotic Animal Diseases under grant no. HSHQDC-16-A-B0006 to JAR.

This work was partially supported by CRIP (Center for Research for Influenza Pathogenesis), a NIAID supportesd Center of Excellence for Influenza Research and Surveillance (CEIRS, contract # HHSN272201400008C) by supplements to NIAID grant U19AI135972 and DoD grant W81XWH-20-1-0270, by the Defense Advanced Research Projects Agency (HR0011-19-2-0020), and by the generous support of the JPB Foundation, the Open Philanthropy Project (research grant 2020-215611 (5384) and anonymous donors to A.G.-S

This work was partially supported by funding to I.M., specifically the Burroughs Wellcome Fund (United States; 1017892); the Chan Zuckerberg Initiative (United States; 2018-191895), the Hirschl Young Investigator fellowship, the NIH UO1 0255-E051 and RO1 0255-B641.

## RESOURCE AVAILABILITY

### Lead Contact

Further information and requests for reagents may be directed to and will be fulfilled by Lead Contact Ivan Marazzi.

### Materials Availability

All unique/stable reagents generated in this study are available from the Lead Contact with a completed Materials Transfer Agreement.

## EXPERIMENTAL MODEL AND SUBJECT DETAILS

### Cells

Human alveolar basal epithelial carcinoma cells (A549, ATCC CCL-185) and monkey kidney epithelial cells (Vero E6, ATCC CRL-1586) were maintained at 37°C and 5% CO_2_ and cultured in Dulbecco’s Modified Eagle’s Medium (DMEM; Gibco) supplemented with 10% fetal bovine serum (FBS; Gibco).

### Viral strains

For infections in A549-ACE2 cells and K18-hACE2 mice, SARS-related coronavirus 2 (SARS-CoV-2), isolate USA-WA1/2020 (NR-52281) was used (Blanco-Melo et al., 2020; Daniloski et al., 2020). For Infections with hamsters in **Figure 3**, isolate HKG/13_P2/2020 (MT835140) was used. For Infections with hamsters in **Figure S2**, isolate USA-WA1/2020 (NR-52281) was used. SARS-CoV-2 was grown in Vero E6 cells in DMEM supplemented with 2% FBS, 4.5 g/L D-glucose, 4 mM L-glutamine, 10 mM non-essential amino acids, 1 mM sodium pyruvate and 10 mM HEPES. Plaque assays were used to determine infectious titers of SARS-CoV-2 by infection of Vero E6 cells in Minimum Essential Media supplemented with 2% FBS, 4 mM L-glutamine, 0.2% BSA, 10 mM HEPES and 0.12% NaHCO3 and 0.7% agar.

## METHOD DETAILS

### Generation of A549-ACE2 Cells

Generation of A549-ACE2 cells was performed as previously described (Blanco-Melo et al., 2020; Daniloski et al., 2020). Briefly, A549 cells were transduced with lentiviral vector pHR-PGK expressing human ACE2 coding sequence. A549 cells were then transduced with the lentivirus in the presence of polybrene (8 μg/ml). Cells were used for downstream assays after 48h post transduction.

### Preparation of siTOP1 sequencing libraries

7.5E4 A549-ACE2 cells were plated in a 24 well dishes. 16 hours post plating, cells were transfected with control, scrambled (siSCR), Top1-targeting (siTOP1) or no siRNA (no siRNA) using Lipofectamine RNAiMax to a final concentration of 50nM. 48hours post transfection, the media was replaced, and fresh media was added to each well. Cells were then mock infected (PBS only) or infected with SARS-CoV-2 at MOI 0.5. 24 hours post infection, media was removed, and cells were lysed in 250ul of Trizol reagent (Thermo Scientific). RNA was then extracted using the Purelink RNA Minikit (Invitrogen) with DNAseI treatment, according to the manufacturer’s recommendations. RNA quality was determined using the RNA 6000 Nano kit and the Eukaryote Total RNA Nano assay on the Agilent 2100 Bioanalyzer System. RNA quantity was determined by Qubit™ RNA HS Assay Kit.

To then prepare RNA-sequencing libraries, 300 ng of RNA was depleted of ribosomal RNA using NEBNext® rRNA Depletion Kit (Human/Mouse/Rat), according to the manusfacturer’s instructions. Libraries were then prepared from rRNA depleted RNA using the NEBNext® Ultra™ II Directional RNA Library Prep Kit for Illumina®, following the manufacturer’s instructions. Final libraries were quantified and sizing was determined using the High Sensitivity DNA Assay reagents and chip in the Agilent 2100 Bioanalyzer System and the Qubit 1X dsDNA HS Assay Kit respectively. Individual libraries were then pooled and sequenced using 75bp paired end on the NextSeq 550 using the NextSeq 500/550 High Output Kit

### ChIP-Seq Library preparation

To prepare ChIP-Sequencing libraries, ~2E5 A549-ACE2 cells were plated into 12 well dishes. Cells were either mock infected (PBS only) or infected with SARS-CoV-2 virus at MOI 0.5. 24 hours post infection, media was removed from the well, and replaced with Fixation buffer (PBS, 2% FBS, 1% Methanol-Free Formaldehye). Cells were fixed at room temperature for 10 min. 2M Glycine was then added to a final concentration of 0.125M, and cells were incubated at room temperature for 5min to quench the reaction. Supernatants were removed from wells, and each well was washed 3 times with cold PBS. Cells were then lysed in the well using 250ul of SDS Lysis Buffer [100mM NaCl, 50mM Tris pH8.0, 5mM EDTA, 0.02% NaN3, 0.5% SDS + 1X Halt Protease and Phosphatase Inhibitor (Thermo Scientific)] and cell lysates were collected in a 1.5ml tube and snap frozen at −80°C. On the day of sonication, lysates were thawed, and diluted with 125ul of Triton Dilution Buffer [100mM Tris pH8.5, 100mM NaCl, 5mM EDTA, 0.02% NaN3, 5% TritonX-100 + 1X Halt Protease and Phosphatase Inhibitor]. Lysates were then sonicated for 5 - 30 sec ON/ 30 sec OFF cycles twice using the Bioruptor Pico. Each sonicated lysate was then pre-cleared using 10ul of Rabbit-IgG Dynabeads for 1 hour, rotating at 4°C. 1ug of anti-H3K27ac antibody was then added to 300ul of pre-cleared lysate. Immunoprecipitation (IP) was performed with overnight rotation at 4°C. To recover IP-complexes, 10ul of Dynabeads M-280 Sheep anti-Rabbit IgG were added to each reaction and tubes were rotated for 2 hours at 4°C. Bead-chromatin complexes were then washed 6 times on a magnet using ice cold RIPA wash buffer [50mM Hepes-KOH pH7.6, 100mM LiCl, 1mM EDTA, 1% NP-40, 0.5% Na-Deoxycholate]. Washed beads were then incubated in 125ul Elution buffer [1% SDS, 0.1M NaHCO3] at 65°C overnight for elution and de-crosslinking. ChIP DNA was then purified using the MinElute PCR purification kit (Qiagen) and quantified using the Qubit 1X dsDNA HS Assay Kit.

ChIP libraries were prepared using the NEBNext^®^ Ultra™ II DNA Library Prep Kit for Illumina following the manufacturer’s recommendations. 1ng of ChIP-DNA was used to prepare each library. ChIP input libraries were prepared by pooling equal amounts of purified sonicated and non-IPed DNA from each sample. 1ng of the pooled ChIP-input DNA was used for library preparation. Libraries were quantified and sizing was determined using the High Sensitivity DNA Assay reagents and chip in the Agilent 2100 Bioanalyzer System and the Qubit 1X dsDNA HS Assay Kit respectively. Individual libraries were then pooled and sequenced 75bp paired end on the NextSeq 550 using the NextSeq 500/550 High Output Kit v2.5.

### Preparation of HiC libraries

In situ Hi-C was performed as described (Heinz 2018) with modifications. The day before infection, 200k A549-ACE2 cells were plated in a 12 well dishes. Cells were either mock-infected (PBS only) or infected with SARS-CoV-2 virus at MOI 0.5. Twenty-four hours post infection, media was removed from the well, and replaced with Fixation buffer (PBS, 2% FBS, 1% Methanol-Free Formaldehyde). Cells were fixed at room temperature for 10 min. 2M Glycine was then added to a final concentration of 0.125M, and cells were incubated at room temperature for 5min to quench the reaction. Supernatants were then removed wells, and each well was washed 2 times with cold PBS. Cells were then lysed in the well using 250 μl of Lysis Buffer [0.5% SDS + Halt Protease and Phosphatase Inhibitors (Thermo Scientific)]. Cell lysates were collected in a 1.5ml tube. 1.5 mU RNAseA (Thermo Scientific) was added to each lysate, and lysates were then incubated at 37°C for 1h. RNAseA treated lysates were then snap frozen and stored in −80°C. After thawing, nuclei were collected at 1500 g for 5 minutes at room temperature. Most of the supernatant was discarded, leaving the nuclei in 10 μl liquid. Samples were resuspended in reaction buffer (25 μl 10% Triton X-100, 25 μl 10x Dpn II buffer, 188 μl water) and rotated at 37°C, 8 RPM for 15 minutes. Chromatin was digested overnight (ON) with either 2 μl (100 U) Dpn II (NEB) (later experiments) at 37°C, rotating overhead with 8 RPM. Nuclei were collected by centrifugation at 1500 g for 5 minutes at room temperature, 225 μl of the supernatant were discarded, leaving the nuclei in 25 μl liquid, and overhangs were filled in with Biotin-14-dATP by adding 75 μl Klenow Master Mix (54.45 μl water, 7.5 μl NEBuffer 2, 0.35 μl 10 mM dCTP, 0.35 μl 10 mM dTTP, 0.35 μl 10 mM dGTP, 7.5 μl 0.4 mM Biotin-14-dATP (Invitrogen), 2 μl 10% Triton X-100, 2.5 μl (12.5 U) Klenow fragment (Enzymatics)) and rotating overhead at RT, 8 RPM for 40 minutes. Reactions were stopped by adding 2.5 μl 0.5 M EDTA and placed on ice. Proximity ligation was performed by transferring the entire reaction into 1.5 ml Eppendorf tubes containing 400 μl ligase mix (322.5 μl water, 40 μl 10x T4 DNA ligase buffer (Enzymatics), 36 μl 10% Triton X-100, 0.5 μl 10 % BSA, 1 μl (600 U) T4 DNA ligase (HC, Enzymatics) and rotating ON at 16°C, 8 RPM. Reactions were stopped with 20 μl 0.5 M EDTA, treated with 1 μl 10 mg/ml DNase-free RNAse A for 15 minutes at 42°C, then 31 μl 5 M NaCl, 29 μl 10 % SDS and 5 μl 20 mg/ml DNase-free proteinase K (Thermo) were added, proteins digested for 1 h at 55°C while shaking at 800 RPM, then crosslinks reversed ON at 65°C. After extraction with 600 μl pH 8-buffered phenol/chloroform/isoamyl alcohol (Ambion) followed by extraction with 600 μl chloroform, DNA was precipitated with 1.5 μl (22.5 μg) Glycoblue (Ambion) and 1400 μl 100% ethanol ON at −20°C. After centrifugation for 20’ at 16000g, 4°C, the DNA pellet was washed twice with 80% ethanol, and the pellet air-dried and dissolved in 131 μl TT (0.05% Tween 20/Tris pH 8). DNA (200 ng) was sheared to 300 bp in 130 μl TT on a Covaris E220 sonicator using the manufacturer’s protocol. Biotinylated DNA was captured on Dynabeads MyOne Streptavidin T1 (Thermo) by combining the sonicated DNA sample (130 μl) with 20 μl Dynabeads that had previously been washed with 1x B&W buffer (2X B&W: 10 mM Tris-HCl pH=7.5, 1 mM EDTA, 2 M NaCl) and had been resuspended in 130 μl 2x B&W containing 0.2% Tween 20. The binding reaction was rotated at RT for 45 minutes, and DNA-bound beads were vigorously washed twice with 150 μl 1x B&W/0.1% Triton-X 100, once with 180 μl TET (0.05% Tween 20, 10 mM Tris pH 8, 1 mM EDTA). Libraries were prepared on-beads using an NEBnext Ultra II DNA library prep kit using half the reagent/reaction volumes given in NEB’s instruction manual and 1.6 pmol Bioo DNA sequencing adapters (Illumina TruSeq-compatible) per reaction. Reactions were stopped by adding 5 μl 0.5 M EDTA, beads collected on a magnet and washed twice with 150 μl 1x B&W/0.1% Triton-X 100, twice with 180 μl TET and resuspended in 20 μl TT (0.05% Tween 20, 10 mM Tris pH 8.0). Libraries were amplified by PCR for 10 cycles (98°C, 30s; 10x [98°C, 10s; 63°C, 20s; 72°C, 30s]; 72°C, 2 min; 4°C, ∞), using 10 μl of the bead suspension in a 50 μl reaction with NEBNext Ultra II Q5 2x master mix (NEB) and 0.5 μM each Solexa 1GA/1GB primers (Solexa 1GA: AATGATACGGCGACCACCGA, Solexa 1GB: CAAGCAGAAGACGGCATACGA). Libraries were precipitated onto magnetic beads by adding 40 μl 20% PEG8000/2.5 M NaCl and 2 μl SpeedBeads (8.9 % PEG final) to 48 μl PCR reaction, thorough mixing by vortexing followed by 10-minute incubation at RT. Beads were collected using a magnet and the supernatant discarded. After washing beads twice by adding 180 μl 80% EtOH, moving the tube strip 6x from side to side of the magnet, collecting beads and discarding the supernatant, beads were air-dried, and DNA eluted by adding 20 μl TT. Libraries were sequenced paired-end for 42 cycles each to a depth of approximately 250 million reads per experiment on an Illumina NextSeq 500 sequencer.

### Preparation of Hamster RNA sequencing libraries

For RNA sequencing analyses in infected hamsters shown in **Figure 3**, infected hamsters that were treated with TPT or vehicle control, were euthanized at days 4 and 6 post infection. Uninfected hamsters were used as controls. After euthanasia, lung left inferior lobe from hamsters were cut into pieces and lysed with RA1 lysis buffer provided with the *NucleoSpin*® *RNA Plus* kit (Macherey-nagel), RNA extraction was performed according the manufacturer’s recommendations, including an on-column genomic DNA digestion step. RNA sequencing library preparation and sequencing were then performed by BGI Genomics

### Hamster Infections

For experiments shown in **Figure 3**, Female Golden Syrian hamster, aged 6-8 week old (~70-100g), were obtained from Laboratory Animal Unit, University of Hong Kong (HKU). All experiments were performed in a Biosafety Level-3 animal facility, LKS Facility of Medicine, HKU. The study has been approved by the Committee on the Use of Live Animals in Teaching and Research, HKU. Virus stock was diluted with Phosphate-buffered saline (PBS) to 2 × 10^4^ PFU/ml. Hamsters were anesthetized with ketamine (150mg/kg) and xylazine (10 mg/mg) and then intranasally inoculation with 50 ul of diluted viruses containing 10^3^ PFU of viruses. For drug treatments, 10mg/kg TPT resuspended in vehicle [5% DMSO + 5% corn oil in PBS] or vehicle alone was administered intraperitoneally to animals on the indicated days post infection.

For experiments shown in **Figure S2**, infection procedures were performed following protocols approved by the Icahn School of Medicine at Mount Sinai Institutional Animal Care and Use Committee (IACUC). Animal studies were carried out in strict accordance with the recommendations in the Guide for the Care and Use of Laboratory Animals of the National Research Council. 7-10 week old (~120-140g) female Golden Syrian hamsters (Charles River) were anesthetized using 90mg/kg Ketamine and 2mg/kg Xylazine by intraperitoneal injection. Once anesthetized, hamsters were intransally infected with 1E5 PFU of SARS-CoV-2 virus re-suspended in 100ul of PBS. Animals were monitored daily for clinical signs of illness and weight loss after infection. For drug treatments, 2mg/kg TPT resuspended in vehicle [4.5% DMSO + 20% Sulfobutylether-β-Cyclodextrin (SBE-β-CD) in PBS] or vehicle alone was administered intraperitoneally to animals on the indicated days post infection.

### K18-hACE2 mice infections

All mice infection procedures were performed following protocols approved by the Icahn School of Medicine at Mount Sinai Institutional Animal Care and Use Committee (IACUC). Animal studies were carried out in strict accordance with the recommendations in the Guide for the Care and Use of Laboratory Animals of the National Research Council. 5-10 week old female B6.Cg-Tg(K18-ACE2)2Prlmn/J (K18-hACE2) mice purchased from Jackson Laboratories (Bar Harbor, ME) were anesthetized by an intraperitoneal injection of 90mg/kg Ketamine and 2 mg/kg xylazine. Once anesthetized, mice were infected with 1E4 PFU of SARS-CoV-2 virus suspended in 30ul of PBS. Mice were monitored daily for clinical signs of illness and weight loss after infection. Animals that reached 75% bodyweight or clinical signs that are irrevocably linked with death were humanely euthanized by intraperitoneal injection of 60mg/kg pentobarbital.

For drug treatments, 2mg/kg Topotecan-hydrochloride (TPT; 14129, Cayman Chemical Company) re-suspended in vehicle [4.5% DMSO + 20% Sulfobutylether-β-Cyclodextrin (SBE-β-CD) in PBS] was administered intraperitoneally to animals on the indicated days post infection.

### Extraction of RNA from Lungs of Infected Hamsters and Mice

Upon euthanasia, the superior lobe of infected hACE2-KI mice or Golden Syrian hamsters were collected for RNA extraction. Lungs were lysed and homogenized in Trizol. RNA extraction was performed using the Purelink RNA Mini Kit with a DNAseI treatment step, according to the manufacturer’s recommendations. cDNA was synthesized from RNA using the High-Capacity cDNA Reverse Transcription Kit (ThermoFisher).

### Histological analysis

For histological slides shown in **Figure 3E to 3H**, Lung left superior lobes of infected Golden Syrian hamster were fixed in 4 % paraformaldehyde and then processed for paraffin embedding. The 4μm tissue sections were stained with haematoxylin and eosin for histopathological examination. Images were with Olympus BX53 semi-motorized fluorescence microscope using cellSens imaging software.

For histological slides shown in **Figure S2D and S2E**, the left lung lobe of infected Golden Syrian hamsters was fixed in 10% formalin for 48 hours. Embedding in paraffin blocks and staining with H&E were conducted by the Biorepository and Pathology Dean’s CoRE at the the Icahn School of Medicine at Mount Sinai. Microscopic sections were analyzed in a blinded fashion by the same pathologist (A.M.). A number was randomly assigned by the investigator to discriminate each section, which was then submitted for analysis. No information about treatments and mouse genotypes was communicated to the pathologist. Lungs were scored by the area involved in broncho-pneumonia.

## QUANTIFICATION AND STATISTICAL ANALYSES

### Mouse Infection Studies

Mice were randomly assigned into treatment groups. Statistical significance between survival curves was calculated using a Log-rank (Mantel-Cox) test using Graphpad Prism 8.0 software. Two tailed Student’s t-tests under the assumption of equal variances between groups were used to compare weight loss in mice from different groups for each day post infection. Data are shown as +-SEM.

### Quantitative qPCR assays

qPCR assays were done with 3-4 biological replicates (3-4 infected animals/condition). Statistical significance in gene expression was estimated with Graphpad Prism 8.0 software, and determined using two-tailed Student’s t test under the assumption of equal variances between groups. Data are shown as +/− SEM

### Illumina Short Read RNA sequencing analyses

After adaptor removal with cutadapt (Martin, 2011) and base-quality trimming to remove 3′read sequences if more than 20 bases with Q <20 were present, paired-end reads were mapped to the SARS-CoV-2 and human (hg38) or hamster (*Mesocricetus auratus*; MesAur1.0) reference genomes with STAR. Gene-count summaries were generated with featureCounts (Liao et al., 2014). A numeric matrix of raw read counts was generated, with genes in rows and samples in columns, and used for differential gene expression analysis with the Bioconductor Limma package (Ritchie et al., 2015) after removing genes with less than 50 total reads across all samples or of less than 200 nucleotides in length. Normalization factors were computed on the filtered data matrix using the weighted trimmed mean of M-values (TMM) method, followed by voom (Law et al., 2014) mean-variance transformation in preparation for Limma linear modeling. To specifically identify the effect of siRNA mediated Top1 depletion on the inflammatory responses to SARS-CoV-2 in A549-ACE2 cells shown in **Figure 2**, we used interaction model (siTOP1_Infected:siTOP_uninfected - siSCR_Infected:siSCR_uninfected or no_siRNA_infected:no_siRNA:uninfected - siSCR_Infected:siSCR_uninfected), that takes into account basal differences between conditions. To identify TPT dependent gene expression changes in infected hamsters showin **in Figure 3**, we performed pairwise contrast between experimental groups (i.e: TPT D4 – DMSO D4; TPT D6 – DMSO D6). Pairwise comparisons were then performed between treatment groups and eBayes adjusted P-values were corrected for multiple testing using the Benjamin-Hochberg (BH) method and used to select genes with significant expression differences (fold change > 1.5, adjusted P value <0.05).

### GSEA analysis for gene signatures in TPT treated hamsters

We identified Top1 inhibitor gene signatures from TPT-treated Syrian hamsters infected with SARS-CoV-2. We defined the up- and down-regulated signatures as genes differentially expressed after 4 or 6 days of treatment (log2|FC| > 1, FDR=10%). We converted hamster genes to available human orthologs using ENSEMBL (Release 101). We downloaded normalized transcript expression from targeted RNA-seq (398 genes) on lung autopsy tissue from COVID19 patients (16 patients, 34 samples), normal lung tissue (6 patients, 17 samples), and lung tissue from from bacterial or viral pneumonia (4 patients, 5 samples) [(Nienhold et al., 2020), GEO accession: GSE151764]. Gene set enrichment analysis (Subramanian et al., 2005) was performed using the R package fgsea (Korotkevich et al., 2019). We used Tukey’s multiple comparison test to identify significant differences in mean normalized enrichment scores.

### ChIP-seq analysis

ChIP-seq datasets were processed and analyzed using an in-house automated pipeline (https://github.com/MarioPujato/NextGenAligner). Briefly, basic quality control for raw sequencing reads was performed using FASTQC (version 0.11.2) (http://www.bioinformatics.babraham.ac.uk/projects/fastqc). Adapter sequences were removed using Trim Galore (version 0.4.2) (https://www.bioinformatics.babraham.ac.uk/projects/trim_galore/), a wrapper script that runs cutadapt (version 1.9.1) to remove the detected adapter sequence from the reads. The quality controlled reads were aligned to the reference human genome (hg19/GRCh37) using bowtie2 (version 2.3.4.1)(Langmead and Salzberg, 2012). Aligned reads were then sorted using samtools (version 1.8)(Li et al., 2009) and duplicate reads were removed using picard (version 1.89) (https://broadinstitute.github.io/picard/). Peaks were called using MACS2 (version 2.1.0) (https://github.com/taoliu/MACS)(Zhang et al., 2008) with the control/input aligned reads as background (callpeak -g hs -q 0.01 –broad -c input/control). ENCODE blacklist regions (Amemiya et al., 2019) were removed using the hg19-blacklist.v2.bed.gz file available at https://github.com/Boyle-Lab/Blacklist/tree/master/lists.

The ChIP-seq experimental design consisted of triplicates experiments for each condition (0hr, 8hr, 24hr infections). PCA analysis indicating strong agreement between experimental replicates and clear separation between conditions (**Figure S1A**) Sequencing reads from replicates were thus combined, and alignment and peak calling was again performed as described above. For differential peak analysis, the union set of all peaks from these three conditions was generated using bedtools (Quinlan and Hall, 2010). For each of the resulting genomic regions, read counts were obtained for all 9 replicates. These read counts used as input to DESeq2 (Love et al., 2014). A fold change cutoff of greater than or equal to 1.5 and an FDR-corrected p-value cutoff of less than or equal to 0.05 were used to identify differential peaks for each pairwise comparison between conditions.

We used the HOMER suite of tools (Heinz et al., 2010), modified to use a log base two scoring system and to include the large set of human motifs contained in the CisBP database (build 2.0)(Lambert et al., 2019) to identify enriched motifs within the sequences of differential and shared ChIP-seq peaks. To minimize redundancy, motifs were grouped into classes using the following procedure. Each human transcription factor was assigned the single best p-value obtained for any of its corresponding motifs. Transcription factors with identical best motifs were merged then into a single class.

### HiC analysis

Hi-C data was processed as described in (Heinz et al., 2018). Briefly, Hi-C reads were trimmed at MboI/DpnII recognition sites (GATC) and aligned to the human genome (GRCh38/hg38) using STAR (Dobin et al., 2013), keeping only read pairs that both map to unique genomic locations for further analysis (MAPQ > 10). All PCR duplicates were also removed. PCA analysis of Hi-C experiments used to define chromatin compartments were performed with HOMER(Lin et al., 2012). For each chromosome, a balanced and distance normalized contact matrix was generated using window size of 50 kb sampled every 25 kb, reporting the ratios of observed to expected contact frequencies for any two regions. The correlation coefficient of the interaction profiles for any two regions across the entire chromosome were then calculated to generate a correlation matrix (also visualized in **Figure 1A**). This matrix was then analyzed using Principal Component Analysis (PCA) from the prcomp function in R (https://www.r-project.org), and the eigenvector loadings for each 25 kb region along the first principal component were assigned to each region (PC1 values). The PC1 values from each chromosome were scaled by their standard deviation to make them more comparable across chromosomes and analysis parameters. For each chromosome, PC1 values are multiplied by −1 if negative PC1 regions are more strongly enriched for active chromatin regions defined by H3K27ac peaks to ensure the positive PC1 values align with the A/permissive compartment (as opposed to the B/inert compartment). chrY was excluded from the PCA analysis due to its small size and high repeat content. Balanced, normalized Hi-C contact maps were generated at 25 kb resolution for visualization (**Figure 1A**). Assignment of PC1 values to Gencode gene promoters and other features was performed using HOMER’s annotatePeaks.pl function using the results from the PCA analysis.

## SUPPLEMENTARY FIGURE LEGENDS

**Figure S1:**
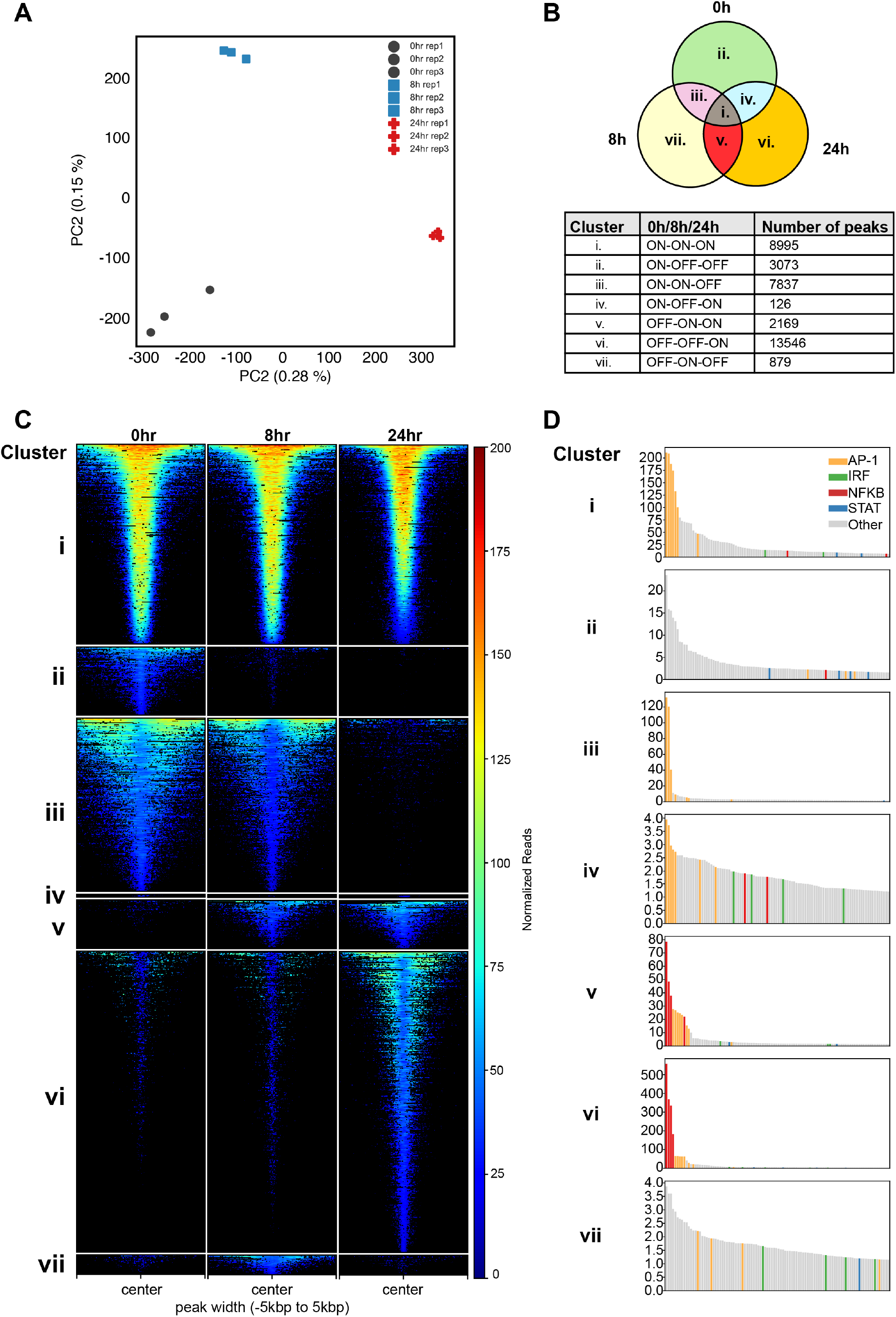
H3K27ac profiles in SARS-CoV-2 infected A549-ACE2 cells. **(A)** Differential H3K27ac across infection time points. H3K27ac ChIP-seq peaks were classified across the infection time course into clusters by their pattern of occurrence (see Methods). **(B)** >Principle Component (PCA) analysis of ChIP-seq experimental replicates. PCA was performed across the genome using the set of peaks identified in each experimental replicate. Percentage of variance expained by the first two components is shown along the axes. PCA was performed using scikit-learn (Pedregosa et al., 2011). **(C)** Venn diagram schematic depicting the seven possible patterns of peak occurence (i-vii), along with the number of peaks observed for each pattern at 0, 8 and 24 hours post infection. **(D)** Heatmap indicating the normalized read count intensity within each peak for each of the three timepoints, for the indicated clusters described in (B). **(E)** Transcription factor binding site motif enrichment for each of the clusters shown in (B) and (C). Motif enrichment was calculated within H3K27ac-marked regions. Bar plots indicate the negative log p-value of enrichment for the top 100 motif classes (see Methods). Bars are colored by motif. AP-1: Yellow; IRF: Green; NFKB: Red; STAT: Blue; Other: Grey

**Figure S2:**
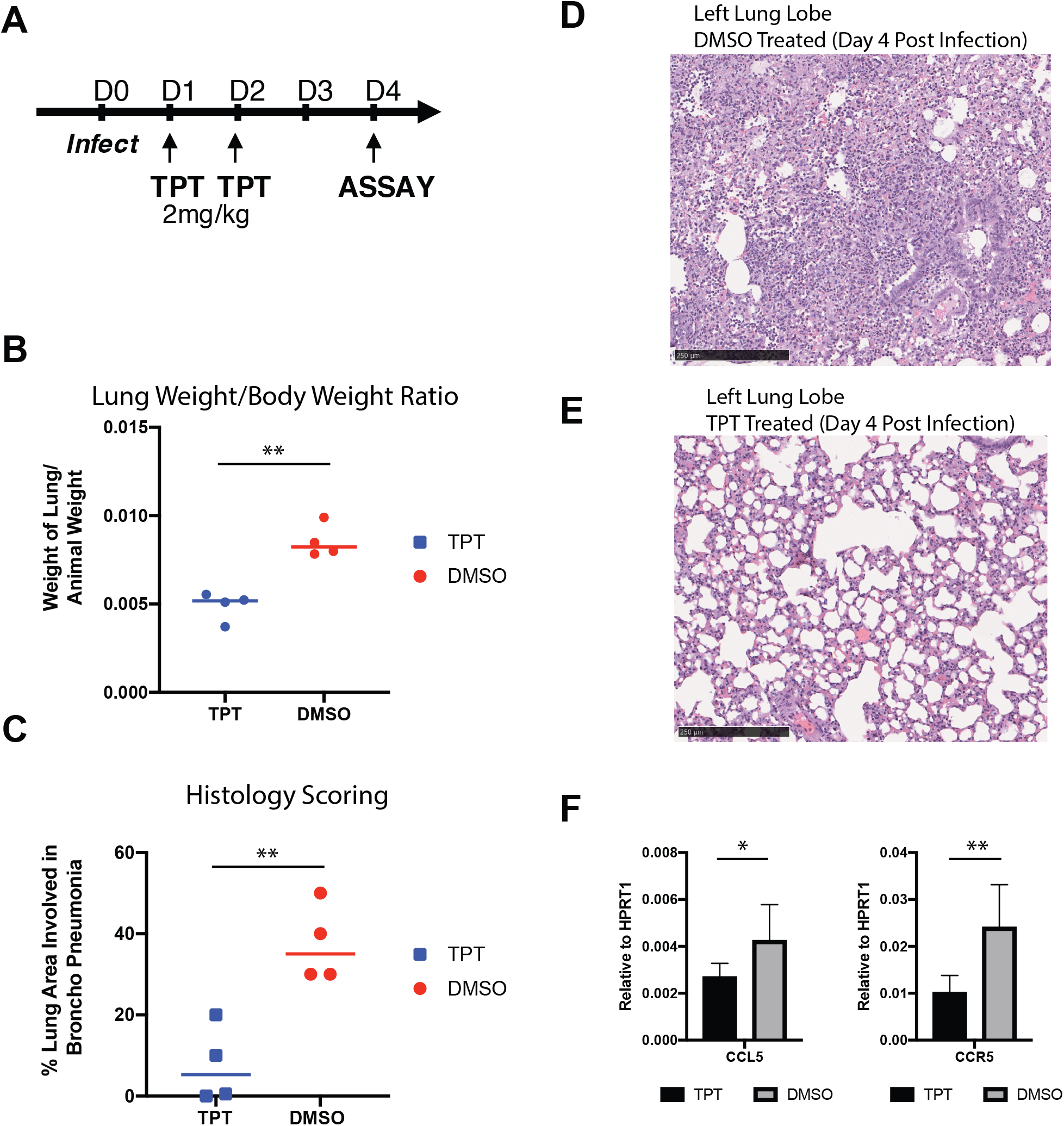
Reduced TPT dosages have similar beneficial effects in SARS-CoV-2 infected hamsters. **(A)** Schematic showing infection and treatment regime in 7-10 week old hamsters. **(B)** Lung weight to body weight ratios of Hamsters infected with 1E4 PFU SARS-CoV-2 at Day 4 post infection, and treated with either DMSO (red) or 2mg/kg TPT (blue). Each dot represents an individual animal, and lines indicate the mean of Lung/Body weight ratios. **(C)** Scatter plots depicting the percent of lung area that is involved in Broncho Pneumonia, as blindly scored by the pathologist (A.M). Each dot represents an individual animal, and the lines indicate the mean. **(D,E)** Representative H&E sections of the left lung lobe of infected hamsters at day 4 post infection, and treated either with DMSO (D) or 2mg/kg TPT (E). Scale bar: 250uM. **(F)** Inflammatory gene expression in DMSO or TPT infected hamsters at day 4 post infection. Bars show the mean and SEM of 4 animals.

## SUPPLEMENTARY TABLE LEGENDS

**Table S1: Table of HiC peak scores and their overlap with H3K27ac peaks (Related to Figure 1)**

Table of processed Hi-C data used to derive panels shown in **Figure 1**. Abbreviations used: IFC: interchromosomal interaction changes: IFC; DLR: distal/local ratio changes; Combo: Replicates combined. Columns T-V: H3K27ac levels in region, at the indicated time points; Columns W-AE: PC1 levels of combined or individual Hi-C replicates; Columns AF-AI: ICF and DLR changes in indicated regions and for the specified contrasts; Column AJ-AQ: Significance values for comparing PC1 values between conditions. Detailed Column descriptions are as follows

**Table S2: H3K27ac ChIP-seq QC summary and differential peak analyses (Related to Figure S1)**

Table S2A: QC analysis and statistics of H3K27ac ChIP-seq data

Table S2B: Full results of H3K27ac differential peak analysis (Contrast 8h vs 0h)

Table S2C: Full results of H3K27ac differential peak analysis (Contrast 24h vs 0h)

**Table S3: H3K27ac ChIP-seq transcription motif analyses summary**

Full results of transcription factor binding site motif enrichment analysis.

**Table S4: Differential expression analysis (RNA-seq) for siRNA treated infected A549-ACE2 cells (Related to Figure 2)**

**Table S4A**: Fold change and associated significance values for genes that are differentially expressed (Fold change > 1.5, p.adj < 0.05) in siTOP1 cells when compared to siSCR treated cells.

**Table S4B**: Same genes shown in Table S4A, but showing the fold changes and associated significance values for in no siRNA treated (no siRNA) cells when compared to siSCR treated cells.

**Table S4C**: Gene lists for all expressed, Top1 dependent and Top1 independent genes. Shown are the log2FC, AveExpr, t-stat, p-values, adjusted p-values and B-stats for the siSCR infected vs siSCR uninfected contrast, used to define infection induced genes. Genes that are induced by infection are indicated by “1” in Column I. Genes that are Top1 dependent and induced by infection are indicated by “1” in Column J. Genes that are Infection induced and Top1 independent are indicated by “1” in Column K.

**Table S5: Gene ontology (pathway) analyses (Related to Figure 2C)**

Gene ontology pathway analysis for downregulated genes listed in Figure 2B and Tables S4.

**Table S6: Blind scores of lung sections from TPT and vehicle (DMSO) treated SARS-CoV-2 Hamsters (Related to Figure S2D and S2E)**

Scoring of H&E stained lung sections taken from hamsters infected with SARS-CoV-2 and treated with vehicle (DMSO) or 2mg/kg TPT at days 1 and 2 post infection. Lung sections were isolated on day 4 post infection.

## REFERENCES

Amemiya, H.M., Kundaje, A., and Boyle, A.P. (2019). The ENCODE Blacklist: Identification of Problematic Regions of the Genome. Sci Rep 9, 9354.

Andre, T., Louvet, C., Maindrault-Goebel, F., Couteau, C., Mabro, M., Lotz, J.P., Gilles-Amar, V., Krulik, M., Carola, E., Izrael, V., et al. (1999). CPT-11 (irinotecan) addition to bimonthly, high-dose leucovorin and bolus and continuous-infusion 5-fluorouracil (FOLFIRI) for pretreated metastatic colorectal cancer. GERCOR. Eur J Cancer 35, 1343–1347.

Banerjee, A.K., Blanco, M.R., Bruce, E.A., Honson, D.D., Chen, L.M., Chow, A., Bhat, P., Ollikainen, N., Quinodoz, S.A., Loney, C., et al. (2020). SARS-CoV-2 Disrupts Splicing, Translation, and Protein Trafficking to Suppress Host Defenses. Cell.

Bao, L., Deng, W., Huang, B., Gao, H., Liu, J., Ren, L., Wei, Q., Yu, P., Xu, Y., Qi, F., et al. (2020). The pathogenicity of SARS-CoV-2 in hACE2 transgenic mice. Nature 583, 830–833.

Bhatraju, P.K., Ghassemieh, B.J., Nichols, M., Kim, R., Jerome, K.R., Nalla, A.K., Greninger, A.L., Pipavath, S., Wurfel, M.M., Evans, L., et al. (2020). Covid-19 in Critically Ill Patients in the Seattle Region - Case Series. N Engl J Med 382, 2012–2022.

Blanco-Melo, D., Nilsson-Payant, B.E., Liu, W.C., Uhl, S., Hoagland, D., Moller, R., Jordan, T.X., Oishi, K., Panis, M., Sachs, D., et al. (2020). Imbalanced Host Response to SARS-CoV-2 Drives Development of COVID-19. Cell 181, 1036–1045 e1039.

Bonev, B., and Cavalli, G. (2016). Organization and function of the 3D genome. Nat Rev Genet 17, 661–678.

Channappanavar, R., Fehr, A.R., Vijay, R., Mack, M., Zhao, J., Meyerholz, D.K., and Perlman, S. (2016). Dysregulated Type I Interferon and Inflammatory Monocyte-Macrophage Responses Cause Lethal Pneumonia in SARS-CoV-Infected Mice. Cell Host Microbe 19, 181–193.

Channappanavar, R., Fehr, A.R., Zheng, J., Wohlford-Lenane, C., Abrahante, J.E., Mack, M., Sompallae, R., McCray, P.B., Jr., Meyerholz, D.K., and Perlman, S. (2019). IFN-I response timing relative to virus replication determines MERS coronavirus infection outcomes. J Clin Invest 129, 3625–3639.

Channappanavar, R., and Perlman, S. (2017). Pathogenic human coronavirus infections: causes and consequences of cytokine storm and immunopathology. Semin Immunopathol 39, 529–539.

Chen, G., Wu, D., Guo, W., Cao, Y., Huang, D., Wang, H., Wang, T., Zhang, X., Chen, H., Yu, H., et al. (2020). Clinical and immunological features of severe and moderate coronavirus disease 2019. J Clin Invest 130, 2620–2629.

Chen, L.F., Lin, Y.T., Gallegos, D.A., Hazlett, M.F., Gomez-Schiavon, M., Yang, M.G., Kalmeta, B., Zhou, A.S., Holtzman, L., Gersbach, C.A., et al. (2019). Enhancer Histone Acetylation Modulates Transcriptional Bursting Dynamics of Neuronal Activity-Inducible Genes. Cell Rep 26, 1174–1188 e1175.

Cheung, C.Y., Poon, L.L., Ng, I.H., Luk, W., Sia, S.F., Wu, M.H., Chan, K.H., Yuen, K.Y., Gordon, S., Guan, Y., et al. (2005). Cytokine responses in severe acute respiratory syndrome coronavirus-infected macrophages in vitro: possible relevance to pathogenesis. J Virol 79, 7819–7826.

Cummings, M.J., Baldwin, M.R., Abrams, D., Jacobson, S.D., Meyer, B.J., Balough, E.M., Aaron, J.G., Claassen, J., Rabbani, L.E., Hastie, J., et al. (2020). Epidemiology, clinical course, and outcomes of critically ill adults with COVID-19 in New York City: a prospective cohort study. Lancet 395, 1763–1770.

Daniloski, Z., Jordan, T.X., Wessels, H.H., Hoagland, D.A., Kasela, S., Legut, M., Maniatis, S., Mimitou, E.P., Lu, L., Geller, E., et al. (2020). Identification of Required Host Factors for SARS-CoV-2 Infection in Human Cells. Cell.

Del Valle, D.M., Kim-Schulze, S., Huang, H.H., Beckmann, N.D., Nirenberg, S., Wang, B., Lavin, Y., Swartz, T.H., Madduri, D., Stock, A., et al. (2020). An inflammatory cytokine signature predicts COVID-19 severity and survival. Nat Med 26, 1636–1643.

Dobin, A., Davis, C.A., Schlesinger, F., Drenkow, J., Zaleski, C., Jha, S., Batut, P., Chaisson, M., and Gingeras, T.R. (2013). STAR: ultrafast universal RNA-seq aligner. Bioinformatics 29, 15–21.

Fukaya, T., Lim, B., and Levine, M. (2016). Enhancer Control of Transcriptional Bursting. Cell 166, 358–368.

Garg, S., Kim, L., Whitaker, M., O’Halloran, A., Cummings, C., Holstein, R., Prill, M., Chai, S.J., Kirley, P.D., Alden, N.B., et al. (2020). Hospitalization Rates and Characteristics of Patients Hospitalized with Laboratory-Confirmed Coronavirus Disease 2019 - COVID-NET, 14 States, March 1-30, 2020. MMWR Morb Mortal Wkly Rep 69, 458–464.

Giamarellos-Bourboulis, E.J., Netea, M.G., Rovina, N., Akinosoglou, K., Antoniadou, A., Antonakos, N., Damoraki, G., Gkavogianni, T., Adami, M.E., Katsaounou, P., et al. (2020). Complex Immune Dysregulation in COVID-19 Patients with Severe Respiratory Failure. Cell Host Microbe 27, 992–1000 e1003.

Grajales-Reyes, G.E., and Colonna, M. (2020). Interferon responses in viral pneumonias. Science 369, 626–627.

Group, W.H.O.R.E.A.f.C.-T.W., Sterne, J.A.C., Murthy, S., Diaz, J.V., Slutsky, A.S., Villar, J., Angus, D.C., Annane, D., Azevedo, L.C.P., Berwanger, O., et al. (2020). Association Between Administration of Systemic Corticosteroids and Mortality Among Critically Ill Patients With COVID-19: A Meta-analysis. JAMA 324, 1330–1341.

Guichard, S., Montazeri, A., Chatelut, E., Hennebelle, I., Bugat, R., and Canal, P. (2001). Schedule-dependent activity of topotecan in OVCAR-3 ovarian carcinoma xenograft: pharmacokinetic and pharmacodynamic evaluation. Clin Cancer Res 7, 3222–3228.

Heinz, S., Benner, C., Spann, N., Bertolino, E., Lin, Y.C., Laslo, P., Cheng, J.X., Murre, C., Singh, H., and Glass, C.K. (2010). Simple combinations of lineage-determining transcription factors prime cis-regulatory elements required for macrophage and B cell identities. Mol Cell 38, 576–589.

Heinz, S., Texari, L., Hayes, M.G.B., Urbanowski, M., Chang, M.W., Givarkes, N., Rialdi, A., White, K.M., Albrecht, R.A., Pache, L., et al. (2018). Transcription Elongation Can Affect Genome 3D Structure. Cell 174, 1522–1536 e1522.

Hermine, O., Mariette, X., Tharaux, P.L., Resche-Rigon, M., Porcher, R., Ravaud, P., and Group, C.-C. (2020). Effect of Tocilizumab vs Usual Care in Adults Hospitalized With COVID-19 and Moderate or Severe Pneumonia: A Randomized Clinical Trial. JAMA Intern Med.

Hildebrand, E.M., and Dekker, J. (2020). Mechanisms and Functions of Chromosome Compartmentalization. Trends Biochem Sci 45, 385–396.

Hnisz, D., Day, D.S., and Young, R.A. (2016). Insulated Neighborhoods: Structural and Functional Units of Mammalian Gene Control. Cell 167, 1188–1200.

Houghton, P.J., Cheshire, P.J., Hallman, J.D., 2nd, Lutz, L., Friedman, H.S., Danks, M.K., and Houghton, J.A. (1995). Efficacy of topoisomerase I inhibitors, topotecan and irinotecan, administered at low dose levels in protracted schedules to mice bearing xenografts of human tumors. Cancer Chemother Pharmacol 36, 393–403.

Huang, C., Wang, Y., Li, X., Ren, L., Zhao, J., Hu, Y., Zhang, L., Fan, G., Xu, J., Gu, X., et al. (2020). Clinical features of patients infected with 2019 novel coronavirus in Wuhan, China. Lancet 395, 497–506.

Kollmannsberger, C., Mross, K., Jakob, A., Kanz, L., and Bokemeyer, C. (1999). Topotecan - A novel topoisomerase I inhibitor: pharmacology and clinical experience. Oncology 56, 1–12.

Korotkevich, G., Sukhov, V., and Sergushichev, A. (2019). Fast gene set enrichment analysis. bioRxiv, 060012.

Lambert, S.A., Yang, A.W.H., Sasse, A., Cowley, G., Albu, M., Caddick, M.X., Morris, Q.D., Weirauch, M.T., and Hughes, T.R. (2019). Similarity regression predicts evolution of transcription factor sequence specificity. Nat Genet 51, 981–989.

Langmead, B., and Salzberg, S.L. (2012). Fast gapped-read alignment with Bowtie 2. Nat Methods 9, 357–359.

Law, C.W., Chen, Y., Shi, W., and Smyth, G.K. (2014). voom: Precision weights unlock linear model analysis tools for RNA-seq read counts. Genome Biol 15, R29.

Lee, S., Channappanavar, R., and Kanneganti, T.D. (2020). Coronaviruses: Innate Immunity, Inflammasome Activation, Inflammatory Cell Death, and Cytokines. Trends Immunol.

Lei, X., Dong, X., Ma, R., Wang, W., Xiao, X., Tian, Z., Wang, C., Wang, Y., Li, L., Ren, L., et al. (2020). Activation and evasion of type I interferon responses by SARS-CoV-2. Nat Commun 11, 3810.

Li, H., Handsaker, B., Wysoker, A., Fennell, T., Ruan, J., Homer, N., Marth, G., Abecasis, G., Durbin, R., and Genome Project Data Processing, S. (2009). The Sequence Alignment/Map format and SAMtools. Bioinformatics 25, 2078–2079.

Liao, Y., Smyth, G.K., and Shi, W. (2014). featureCounts: an efficient general purpose program for assigning sequence reads to genomic features. Bioinformatics 30, 923–930.

Lin, Y.C., Benner, C., Mansson, R., Heinz, S., Miyazaki, K., Miyazaki, M., Chandra, V., Bossen, C., Glass, C.K., and Murre, C. (2012). Global changes in the nuclear positioning of genes and intra- and interdomain genomic interactions that orchestrate B cell fate. Nat Immunol 13, 1196–1204.

Liu, X., Hong, T., Parameswaran, S., Ernst, K., Marazzi, I., Weirauch, M.T., and Fuxman Bass, J.I. (2020). Human Virus Transcriptional Regulators. Cell 182, 24–37.

Livingston, E., and Bucher, K. (2020). Coronavirus Disease 2019 (COVID-19) in Italy. JAMA 323, 1335.

Love, M.I., Huber, W., and Anders, S. (2014). Moderated estimation of fold change and dispersion for RNA-seq data with DESeq2. Genome Biol 15, 550.

Lucas, C., Wong, P., Klein, J., Castro, T.B.R., Silva, J., Sundaram, M., Ellingson, M.K., Mao, T., Oh, J.E., Israelow, B., et al. (2020). Longitudinal analyses reveal immunological misfiring in severe COVID-19. Nature 584, 463–469.

Marazzi, I., Greenbaum, B.D., Low, D.H.P., and Guccione, E. (2018). Chromatin dependencies in cancer and inflammation. Nat Rev Mol Cell Biol 19, 245–261.

Marazzi, I., Ho, J.S., Kim, J., Manicassamy, B., Dewell, S., Albrecht, R.A., Seibert, C.W., Schaefer, U., Jeffrey, K.L., Prinjha, R.K., et al. (2012). Suppression of the antiviral response by an influenza histone mimic. Nature 483, 428–433.

Martin, M. (2011). Cutadapt removes adapter sequences from high-throughput sequencing reads. 2011 17, 3.

Mathijssen, R.H., van Alphen, R.J., Verweij, J., Loos, W.J., Nooter, K., Stoter, G., and Sparreboom, A. (2001). Clinical pharmacokinetics and metabolism of irinotecan (CPT-11). Clin Cancer Res 7, 2182–2194.

Merad, M., and Martin, J.C. (2020). Pathological inflammation in patients with COVID-19: a key role for monocytes and macrophages. Nat Rev Immunol 20, 355–362.

Miller, M.S., Rialdi, A., Ho, J.S., Tilove, M., Martinez-Gil, L., Moshkina, N.P., Peralta, Z., Noel, J., Melegari, C., Maestre, A.M., et al. (2015). Senataxin suppresses the antiviral transcriptional response and controls viral biogenesis. Nat Immunol 16, 485–494.

Moore, J.B., and June, C.H. (2020). Cytokine release syndrome in severe COVID-19. Science 368, 473–474.

Munoz-Fontela, C., Dowling, W.E., Funnell, S.G.P., Gsell, P.S., Riveros-Balta, A.X., Albrecht, R.A., Andersen, H., Baric, R.S., Carroll, M.W., Cavaleri, M., et al. (2020). Animal models for COVID-19. Nature 586, 509–515.

Nemati, F., Daniel, C., Arvelo, F., Legrier, M.E., Froget, B., Livartowski, A., Assayag, F., Bourgeois, Y., Poupon, M.F., and Decaudin, D. (2010). Clinical relevance of human cancer xenografts as a tool for preclinical assessment: example of in-vivo evaluation of topotecan-based chemotherapy in a panel of human small-cell lung cancer xenografts. Anticancer Drugs 21, 25–32.

Nicodeme, E., Jeffrey, K.L., Schaefer, U., Beinke, S., Dewell, S., Chung, C.W., Chandwani, R., Marazzi, I., Wilson, P., Coste, H., et al. (2010). Suppression of inflammation by a synthetic histone mimic. Nature 468, 1119–1123.

Nienhold, R., Ciani, Y., Koelzer, V.H., Tzankov, A., Haslbauer, J.D., Menter, T., Schwab, N., Henkel, M., Frank, A., Zsikla, V., et al. (2020). Two distinct immunopathological profiles in autopsy lungs of COVID-19. Nat Commun 11, 5086.

O’Driscoll, M., Dos Santos, G.R., Wang, L., Cummings, D.A.T., Azman, A.S., Paireau, J., Fontanet, A., Cauchemez, S., and Salje, H. (2020). Age-specific mortality and immunity patterns of SARS-CoV-2. Nature.

Oran, D.P., and Topol, E.J. (2020). Prevalence of Asymptomatic SARS-CoV-2 Infection: A Narrative Review. Ann Intern Med 173, 362–367.

Pedregosa, F., Varoquaux, G., Gramfort, A., Michel, V., Thirion, B., Grisel, O., Blondel, M., Prettenhofer, P., Weiss, R., Dubourg, V., et al. (2011). Scikit-learn: Machine Learning in Python. Journal of Machine Learning Research 12, 2825–2830.

Qin, C., Zhou, L., Hu, Z., Zhang, S., Yang, S., Tao, Y., Xie, C., Ma, K., Shang, K., Wang, W., et al. (2020). Dysregulation of Immune Response in Patients With Coronavirus 2019 (COVID-19) in Wuhan, China. Clin Infect Dis 71, 762–768.

Quinlan, A.R., and Hall, I.M. (2010). BEDTools: a flexible suite of utilities for comparing genomic features. Bioinformatics 26, 841–842.

Rao, S.S.P., Huang, S.C., Glenn St Hilaire, B., Engreitz, J.M., Perez, E.M., Kieffer-Kwon, K.R., Sanborn, A.L., Johnstone, S.E., Bascom, G.D., Bochkov, I.D., et al. (2017). Cohesin Loss Eliminates All Loop Domains. Cell 171, 305–320 e324.

Rialdi, A., Campisi, L., Zhao, N., Lagda, A.C., Pietzsch, C., Ho, J.S.Y., Martinez-Gil, L., Fenouil, R., Chen, X., Edwards, M., et al. (2016). Topoisomerase 1 inhibition suppresses inflammatory genes and protects from death by inflammation. Science 352, aad7993.

Ritchie, M.E., Phipson, B., Wu, D., Hu, Y., Law, C.W., Shi, W., and Smyth, G.K. (2015). limma powers differential expression analyses for RNA-sequencing and microarray studies. Nucleic Acids Res 43, e47.

Rowinsky, E.K., Grochow, L.B., Hendricks, C.B., Ettinger, D.S., Forastiere, A.A., Hurowitz, L.A., McGuire, W.P., Sartorius, S.E., Lubejko, B.G., Kaufmann, S.H., et al. (1992). Phase I and pharmacologic study of topotecan: a novel topoisomerase I inhibitor. J Clin Oncol 10, 647–656.

Salvarani, C., Dolci, G., Massari, M., Merlo, D.F., Cavuto, S., Savoldi, L., Bruzzi, P., Boni, F., Braglia, L., Turra, C., et al. (2020). Effect of Tocilizumab vs Standard Care on Clinical Worsening in Patients Hospitalized With COVID-19 Pneumonia: A Randomized Clinical Trial. JAMA Intern Med.

Schwarzer, W., Abdennur, N., Goloborodko, A., Pekowska, A., Fudenberg, G., Loe-Mie, Y., Fonseca, N.A., Huber, W., Haering, C.H., Mirny, L., et al. (2017). Two independent modes of chromatin organization revealed by cohesin removal. Nature 551, 51–56.

Senecal, A., Munsky, B., Proux, F., Ly, N., Braye, F.E., Zimmer, C., Mueller, F., and Darzacq, X. (2014). Transcription factors modulate c-Fos transcriptional bursts. Cell Rep 8, 75–83.

Siddiqi, H.K., and Mehra, M.R. (2020). COVID-19 illness in native and immunosuppressed states: A clinical-therapeutic staging proposal. J Heart Lung Transplant 39, 405–407.

Slutsky, A.S., and Ranieri, V.M. (2013). Ventilator-induced lung injury. N Engl J Med 369, 2126–2136.

Stokes, E.K., Zambrano, L.D., Anderson, K.N., Marder, E.P., Raz, K.M., El Burai Felix, S., Tie, Y., and Fullerton, K.E. (2020). Coronavirus Disease 2019 Case Surveillance - United States, January 22-May 30, 2020. MMWR Morb Mortal Wkly Rep 69, 759–765.

Subramanian, A., Tamayo, P., Mootha, V.K., Mukherjee, S., Ebert, B.L., Gillette, M.A., Paulovich, A., Pomeroy, S.L., Golub, T.R., Lander, E.S., et al. (2005). Gene set enrichment analysis: a knowledge-based approach for interpreting genome-wide expression profiles. Proc Natl Acad Sci U S A 102, 15545–15550.

Tang, Y., Liu, J., Zhang, D., Xu, Z., Ji, J., and Wen, C. (2020). Cytokine Storm in COVID-19: The Current Evidence and Treatment Strategies. Front Immunol 11, 1708.

von Pawel, J., Schiller, J.H., Shepherd, F.A., Fields, S.Z., Kleisbauer, J.P., Chrysson, N.G., Stewart, D.J., Clark, P.I., Palmer, M.C., Depierre, A., et al. (1999). Topotecan versus cyclophosphamide, doxorubicin, and vincristine for the treatment of recurrent small-cell lung cancer. J Clin Oncol 17, 658–667.

Wang, D., Hu, B., Hu, C., Zhu, F., Liu, X., Zhang, J., Wang, B., Xiang, H., Cheng, Z., Xiong, Y., et al. (2020a). Clinical Characteristics of 138 Hospitalized Patients With 2019 Novel Coronavirus-Infected Pneumonia in Wuhan, China. JAMA 323, 1061–1069.

Wang, D., Li, R., Wang, J., Jiang, Q., Gao, C., Yang, J., Ge, L., and Hu, Q. (2020b). Correlation analysis between disease severity and clinical and biochemical characteristics of 143 cases of COVID-19 in Wuhan, China: a descriptive study. BMC Infect Dis 20, 519.

Winkler, E.S., Bailey, A.L., Kafai, N.M., Nair, S., McCune, B.T., Yu, J., Fox, J.M., Chen, R.E., Earnest, J.T., Keeler, S.P., et al. (2020). SARS-CoV-2 infection of human ACE2-transgenic mice causes severe lung inflammation and impaired function. Nat Immunol 21, 1327–1335.

Wong, C.K., Lam, C.W., Wu, A.K., Ip, W.K., Lee, N.L., Chan, I.H., Lit, L.C., Hui, D.S., Chan, M.H., Chung, S.S., et al. (2004). Plasma inflammatory cytokines and chemokines in severe acute respiratory syndrome. Clin Exp Immunol 136, 95–103.

Wu, C., Chen, X., Cai, Y., Xia, J., Zhou, X., Xu, S., Huang, H., Zhang, L., Zhou, X., Du, C., et al. (2020). Risk Factors Associated With Acute Respiratory Distress Syndrome and Death in Patients With Coronavirus Disease 2019 Pneumonia in Wuhan, China. JAMA Intern Med 180, 934–943.

Wu, Z., and McGoogan, J.M. (2020). Characteristics of and Important Lessons From the Coronavirus Disease 2019 (COVID-19) Outbreak in China: Summary of a Report of 72314 Cases From the Chinese Center for Disease Control and Prevention. JAMA 323, 1239–1242.

Yang, X., Yu, Y., Xu, J., Shu, H., Xia, J., Liu, H., Wu, Y., Zhang, L., Yu, Z., Fang, M., et al. (2020a). Clinical course and outcomes of critically ill patients with SARS-CoV-2 pneumonia in Wuhan, China: a single-centered, retrospective, observational study. Lancet Respir Med 8, 475–481.

Yang, Y., Shen, C., Li, J., Yuan, J., Wei, J., Huang, F., Wang, F., Li, G., Li, Y., Xing, L., et al. (2020b). Plasma IP-10 and MCP-3 levels are highly associated with disease severity and predict the progression of COVID-19. J Allergy Clin Immunol 146, 119–127 e114.

Zabidi, M.A., Arnold, C.D., Schernhuber, K., Pagani, M., Rath, M., Frank, O., and Stark, A. (2015). Enhancer-core-promoter specificity separates developmental and housekeeping gene regulation. Nature 518, 556–559.

Zhang, Y., Liu, T., Meyer, C.A., Eeckhoute, J., Johnson, D.S., Bernstein, B.E., Nusbaum, C., Myers, R.M., Brown, M., Li, W., et al. (2008). Model-based analysis of ChIP-Seq (MACS). Genome Biol 9, R137.

Zhou, F., Yu, T., Du, R., Fan, G., Liu, Y., Liu, Z., Xiang, J., Wang, Y., Song, B., Gu, X., et al. (2020a). Clinical course and risk factors for mortality of adult inpatients with COVID-19 in Wuhan, China: a retrospective cohort study. Lancet 395, 1054–1062.

Zhou, Y., Fu, B., Zheng, X., Wang, D., Zhao, C., Qi, Y., Sun, R., Tian, Z., Xu, X., and Wei, H. (2020b). Pathogenic T-cells and inflammatory monocytes incite inflammatory storms in severe COVID-19 patients. National Science Review 7, 998–1002.

Zhu, Z., Lian, X., Su, X., Wu, W., Marraro, G.A., and Zeng, Y. (2020). From SARS and MERS to COVID-19: a brief summary and comparison of severe acute respiratory infections caused by three highly pathogenic human coronaviruses. Respir Res 21, 224.

